# *Brucella* periplasmic protein EipB is a molecular determinant of cell envelope integrity and virulence

**DOI:** 10.1101/551135

**Authors:** Julien Herrou, Jonathan W. Willett, Aretha Fiebig, Daniel M. Czyż, Jason X. Cheng, Eveline Ultee, Ariane Briegel, Lance Bigelow, Gyorgy Babnigg, Youngchang Kim, Sean Crosson

**Affiliations:** Department of Biochemistry and Molecular Biology, University of Chicago, Chicago, Illinois, USA; Department of Microbiology and Cell Science, University of Florida, Gainesville, Florida, USA; Department of Pathology, The University of Chicago, Chicago, Illinois, USA; Department of Biology, Universiteit Leiden, Leiden, Netherlands; Biosciences Division, Argonne National Laboratory, Argonne, Illinois, USA

**Author notes:** To whom correspondence should be addressed: Sean Crosson,. Contributed equally to this work (Julien Herrou, Jonathan W. Willett). Current location: Laboratoire de Chimie Bactérienne, UMR7283, Institut de Microbiologie de la Méditerranée, CNRS, Marseille, France.

**Keywords:** TPR, DUF1849, *Alphaproteobacteria*, *Brucella*, cell envelope, stress response, PF08904

## Abstract

The Gram-negative cell envelope is a remarkable structure with core components that include an inner membrane, an outer membrane, and a peptidoglycan layer in the periplasmic space between. Multiple molecular systems function to maintain integrity of this essential barrier between the interior of the cell and its surrounding environment. We show that a conserved DUF1849-family protein, EipB, is secreted to the periplasmic space of *Brucella*, a monophyletic group of intracellular pathogens. In the periplasm, EipB folds into an unusual fourteen-stranded β-spiral structure that resembles the LolA and LolB lipoprotein delivery system, though the overall fold of EipB is distinct from LolA/LolB. Deletion of *eipB* results in defects in *Brucella* cell envelope integrity *in vitro* and in maintenance of spleen colonization in a mouse model of *B. abortus* infection. Transposon disruption of *ttpA*, which encodes a periplasmic protein containing tetratricopeptide repeats, is synthetically lethal with *eipB* deletion. *ttpA* is a reported virulence determinant in *Brucella*, and our studies of *ttpA* deletion and overexpression strains provide evidence that this gene also contributes to cell envelope function. We conclude that *eipB* and *ttpA* function in the *Brucella* periplasmic space to maintain cell envelope integrity, which facilitates survival in a mammalian host.

**Importance:** *Brucella* species cause brucellosis, a global zoonosis. A gene encoding a conserved DUF1849-family protein, which we have named EipB, is present in all sequenced *Brucella* and several other genera in the class *Alphaproteobacteria*. This manuscript provides the first functional and structural characterization of a DUF1849 protein. We show that EipB is secreted to the periplasm where it forms a spiral-shaped antiparallel-β protein that is a determinant of cell envelope integrity *in vitro* and virulence in an animal model of disease. *eipB* genetically interacts with *ttpA*, which also encodes a periplasmic protein. We propose that EipB and TtpA function as part of a system required for cell envelope homeostasis in select *Alphaproteobacteria*.

## Introduction

*Brucella* spp. are the causative agents of brucellosis, which afflicts wildlife and livestock on a global scale and can occur in humans through contact with infected animals or animal products (1, 2). These intracellular pathogens are members of the class *Alphaproteobacteria*, a group of Gram-negative species that exhibit tremendous diversity in metabolic capacity, cell morphology, and ecological niches (3). In their mammalian hosts, *Brucella* cells must contend with the host immune system (4) and adapt to stresses including oxidative assault from immune cells, acidic pH in the phagosomal compartment, and nutrient shifts during intracellular trafficking (5). Molecular components of the cell envelope play a key role in the ability of *Brucella* spp. to survive these stresses and to replicate in the intracellular niche (6, 7). As part of a systematic experimental survey of conserved *Alphaproteobacterial* protein **<u>d</u>**omains of **<u>u</u>**nknown **<u>f</u>**unction (DUFs), we recently described **<u>e</u>**nvelope **<u>i</u>**ntegrity **<u>p</u>**rotein **<u>A</u>** (EipA). This periplasmic protein confers resistance to cell envelope stressors and determines *B. abortus* virulence in a mouse model of infection (8). In this study, we report a functional and structural analysis of envelope integrity **<u>p</u>**rotein **<u>B</u>** (EipB), a member of the uncharacterized gene family DUF1849.

DUF1849 (Pfam: PF08904, (9)) is widespread among the *Rhizobiales, Rhodospirillales* and *Rhodobacterales* (Figure 1). To our knowledge, no functional data have been reported for this gene family other than results from a recent multi-species Tn-seq study that showed stress sensitivity in *Sinorhizobium meliloti* DUF1849 (locus *SMc02102*) mutant strains (10). Here we show that the *Brucella* DUF1849 protein, EipB (locus tag *bab1_1186;* RefSeq locus BAB_RS21600), is a 280-residue periplasmic protein that folds into a 14-stranded, open **β**-barrel structure containing a conserved disulfide bond. We term this novel barrel structure a **β**-spiral and show that it resembles the lipoprotein chaperone LolB, though its overall fold is distinct. Replication and survival of a *B. abortus* strain in which we deleted *eipB* was attenuated in a mouse infection model, and deletion of *eipB* in both *B. abortus* and *Brucella ovis* enhanced sensitivity to compounds that affect the integrity of the cell envelope. We have further shown that *B. abortus eipB* deletion is synthetically lethal with transposon disruption of gene locus *bab1_0430*, which encodes a periplasmic **<u>t</u>**etra**<u>t</u>**ricopeptide-repeat (TPR) containing-**<u>p</u>**rotein that we have named TtpA. The *Brucella melitensis* ortholog of TtpA (locus tag BMEI1531) has been previously described as a molecular determinant of mouse spleen colonization (11), while a *Rhizobium leguminosarum* TtpA homolog (locus tag RL0936) is required for proper cell envelope function (12). We propose that TtpA and EipB coordinately function in the *Brucella* periplasm to ensure cell envelope integrity and to enable cell survival in the mammalian host niche.

**Figure 1:**
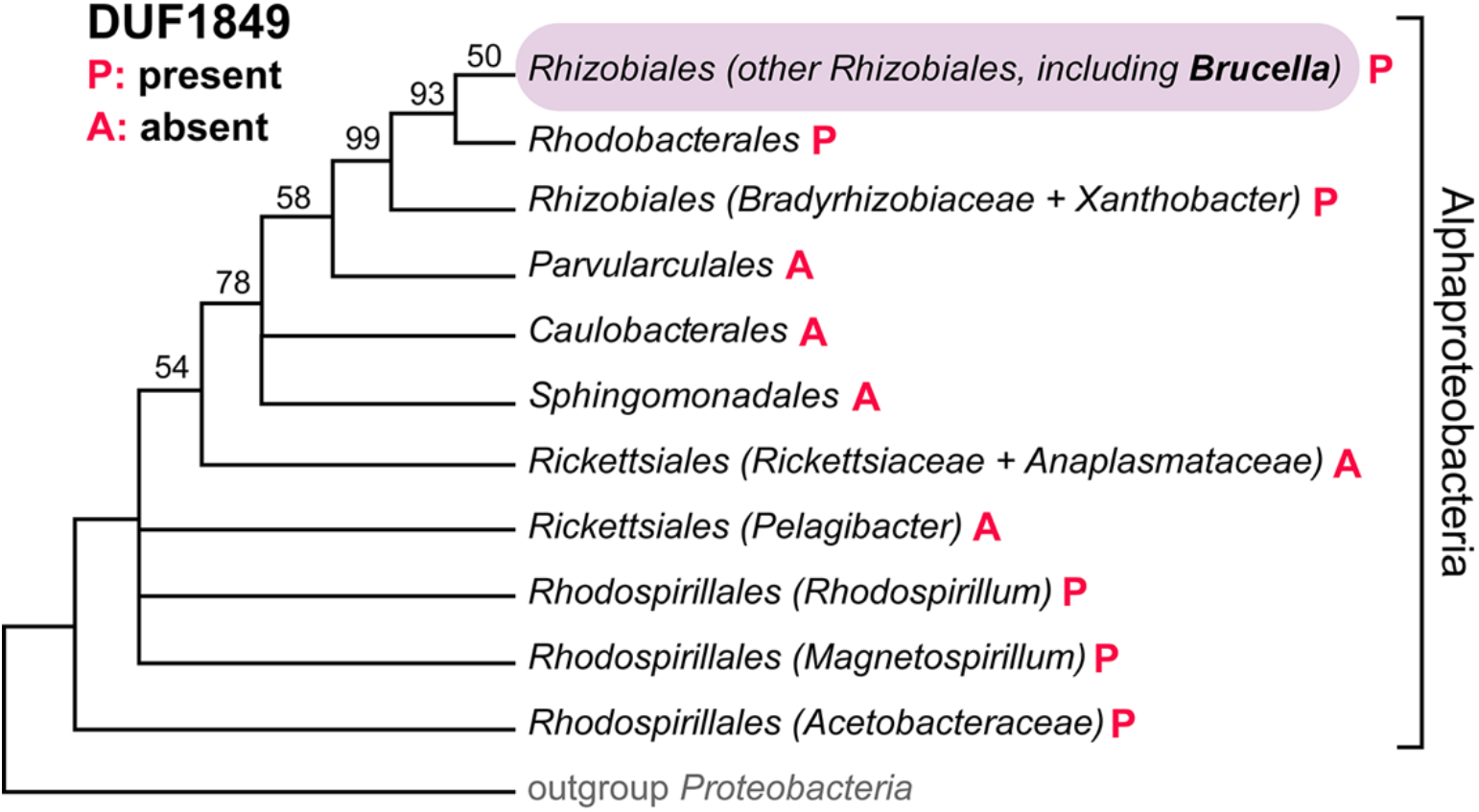
The DUF1849 sequence family is restricted to *Alphaproteobacteria*. Bayesian phylogenetic tree showing the distribution of DUF1849 genes in different orders within the class *Alphaproteobacteria* (P: present, A: absent). Bayesian support values are shown when <100%; nodes were collapsed when support was <50%; adapted from Williams and colleagues (50). In *Brucella abortus* (order *Rhizobiales*), DUF1849 is encoded by gene locus *bab1_1186* (i.e. *eipB*).

## Results

### *B. abortus eipB* is required for maintenance of mouse spleen colonization

As part of a screen to evaluate the role of conserved *Alphaproteobacterial* genes of unknown function in *B. abortus* infection biology, we infected THP-1 macrophage-like cells with wild-type *B. abortus*, an *eipB* deletion strain (Δ*eipB*), and a genetically complemented Δ*eipB* strain. Infected macrophages were lysed and colony forming units (CFU) were enumerated on tryptic soy agar plates (TSA) at 1, 24 and 48 hours post-infection. We observed no significant differences between strains at 1, 24 or 48 hours post-infection, indicating that *eipB* was not required for entry, replication or intracellular survival *in vitro* (Figure 2A).

**Figure 2:**
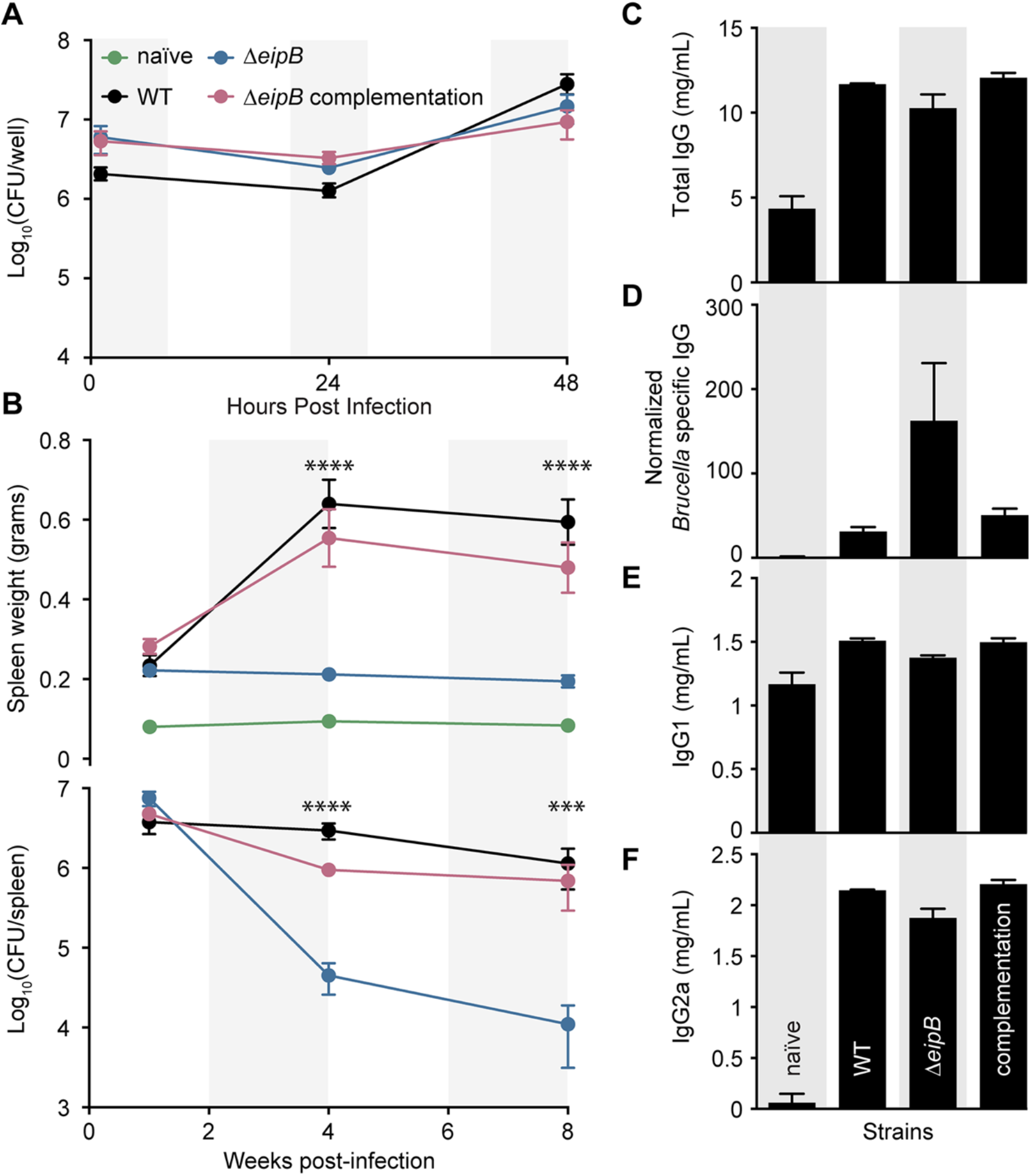
*eipB* is a genetic determinant of *B. abortus* virulence. A) *In vitro* macrophage infection assay: infection of THP-1 cells with wild-type *B. abortus* 2308 (black line), Δ*eipB* (blue line) and the *eipB* complementation strain (pink line). The number of *B. abortus* CFUs recovered from the THP-1 cells at 1, 24, and 48 hours post infection is plotted. Each data point (n= 3 per strain) is the mean ± the standard error of the mean. B) *In vivo* mouse infection assay: female BALB/c mice were injected intraperitoneally with wild-type, Δ*eipB*, or Δ*eipB*-complementation strains. Spleen weights (upper graph) and bacterial burden (lower graph) were measured at 1, 4, and 8 weeks post-infection. Graphs represent data from uninfected, naïve mice (in green) or mice infected with wild-type (black), Δ*eipB* (blue), or complementation (pink) strains. Data presented are the mean ± the standard error of the mean; n= 5 mice per strain per time point. One-way ANOVA followed by Dunnett’s post test (to wild-type) supports the conclusion that spleens infected with the *eipB* deletion strain were significantly smaller at 4 (****, p<0.0001) and 8 weeks (****, p<0.0001) and had fewer CFU than wild-type at 4 (****, p<0.0001) and 8 weeks (***, p<0.0007). C-F) Antibody quantification in mouse serum harvested at 8 weeks post-infection from naïve control mice or mice infected with wild-type, Δ*eipB*, or complementation strains. Amounts of total IgG at 8 weeks (C), Brucella-specific IgG (D), IgG1 (E), and IgG2a (F) were determined by ELISA. Each data point (naïve: n= 3, WT: n= 2, Δ*eipB* and complementation: n= 4) is the mean ± the standard error of the mean.

We further evaluated the role of *eipB* in a BALB/c mouse infection model. Mice infected with Δ*eipB* had no significant difference in spleen weight or bacterial load compared to mice infected with wild-type *B. abortus* strain 2308 at one-week post-infection (Figure 2B). However, at 4- and 8-weeks post-infection, mice infected with the wild-type or the complemented *eipB* deletion strains had pronounced splenomegaly and a bacterial load of approximately 5 x 10^6^ CFU/spleen. In contrast, mice infected with Δ*eipB* had smaller spleens with approximately 2 orders fewer bacteria (~1 x 10^4^ CFU/spleen) (Figure 2B). We conclude that *eipB* is not required for initial spleen colonization but is necessary for full virulence and persistence in the spleen over an 8-week time course.

To assess the pathology of mice infected with wild-type and Δ*eipB* strains, we harvested spleens at 8 weeks post-infection and fixed, mounted, and subjected the samples to hematoxylin and eosin (H&E) staining (Figure S1). Compared to naïve (uninfected) mice (Figure S1A), we observed higher extramedullary hematopoiesis, histiocytic proliferation, granulomas, and the presence of *Brucella* immunoreactivities in spleens of mice infected with wild-type *B. abortus* 2308 and the genetically-complemented mutant strain (Figure S1B and D). Both wild-type and the complemented strain caused spleen inflammation with a reduced white to red pulp ratio as a result of lymphoid follicle disruption and red pulp expansion, which typically correlates with infiltration of inflammatory cells; these spleens also had increased marginal zones (Figure S1B and D). As expected from the CFU enumeration data, mice infected with Δ*eipB* had reduced pathologic features: there was minimal change in white to red pulp ratio, and a minimal increase in marginal zones (Figure S1C). There was no evidence of extramedullary hematopoiesis in mice infected with Δ*eipB*, though histiocytic proliferation was mildly increased. Granulomas and *Brucella* immunoreactivities were rare in Δ*eipB* (Figure S1C). These results are consistent with a model in which *eipB* is required for full *B. abortus* virulence in a mouse model of infection. A summary of spleen pathology scores is presented in Table S1.

We further measured antibody responses in mice infected with Δ*eipB* and wild-type strains. Serum levels of total IgG, Brucella-specific IgG, subclass IgG1, and subclass IgG2a were measured by enzyme-linked immunosorbent assays (ELISA) (Figure 2C-F). Antibody subclasses IgG2a and IgG1 were measured as markers of T helper 1 (Th1)- and Th2-specific immune responses, respectively. At 8 weeks post-infection, total serum IgG was higher in all infected mice relative to the uninfected control (Figure 2C). The level of Brucella-specific IgG was approximately 5 times higher in Δ*eipB*-infected mice than in mice infected with wild-type or the complemented mutant strain (Figure 2D). Uninfected mice and mice infected with wild-type, Δ*eipB* and the Δ*eipB*-complemented strain showed no significant difference in IgG1 levels after 8 weeks (Figure 2E). All infected mice had highly increased levels of IgG2a at 8 weeks post infection relative to naïve mice, though there was no difference between *B. abortus* strains (Figure 2F). We conclude that Δ*eipB* infection results in production of more *B. abortus-specific* antibodies than wild-type. Subclasses IgG1 and IgG2a do not apparently account for the higher levels of these specific antibodies. Large induction of IgG2a by all *B. abortus* strains is consistent with the known ability of *B. abortus* to promote a strong Th1 response (13, 14). However, Δ*eipB* does not induce a more robust Th1 response than wild-type based on our IgG2a measurements. We did not test whether antibodies contribute to clearance of the Δ*eipB* strain. Enhanced *Brucella*-specific antibody production may simply be a consequence of antigen release triggered by host clearance of Δ*eipB* by other immune mechanisms.

### The Δ*eipB* strain is sensitive to cell envelope stressors

To test whether reduced virulence of Δ*eipB* correlates with an increased sensitivity to stress *in vitro*, we evaluated *B. abortus* Δ*eipB* growth on TSA plates supplemented with known cell membrane/envelope stressors including EDTA, ampicillin and deoxycholate. Δ*eipB* had 1.5 to 3 orders fewer CFUs compared to wild-type when titered on TSA plates containing these compounds. All phenotypes were complemented by restoring the Δ*eipB* locus to wild-type (Figure 3A). Together, these data provide evidence that *eipB* contributes to resistance to compounds that compromise the integrity of the *B. abortus* cell membrane/envelope.

**Figure 3:**
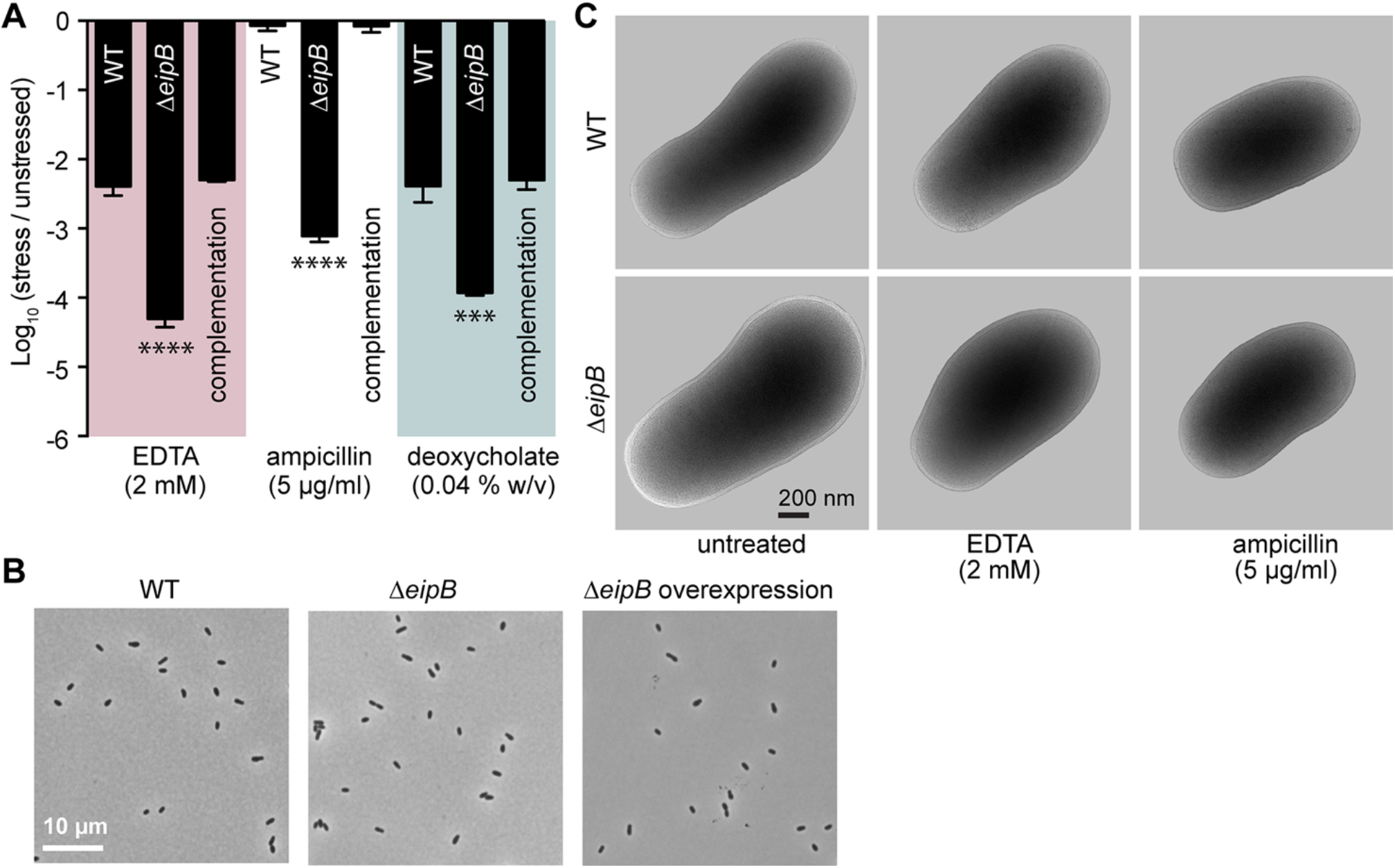
Assessing the effect of cell envelope stressors on *B. abortus* Δ*eipB* growth and survival. A) Envelope stress survival assays. Serially diluted cultures of *B. abortus* wild-type, Δ*eipB*, and complementation strains were spotted on plain TSA plates or TSA plates containing EDTA (2 mM), deoxycholate (0.04% w/v), or ampicillin (5 μg/ml). After 3 to 5 days of growth at 37°C / 5% CO2, CFUs for each condition were enumerated and plotted. This experiment was repeated four times; each data point is the mean ± the standard error of the mean. One-way ANOVA followed by Dunnett’s post test (to wild-type) supports the conclusion that the *eipB* deletion strain had significantly fewer CFU than wild-type in presence of EDTA (****, p<0.0001), ampicillin (****, p<0.0001), and deoxycholate (***, p<0.0003). B) Light micrograph of *B. abortus* wild-type (left), Δ*eipB* (middle) and overexpression (right; induced with 5 mM IPTG) liquid cultures grown overnight in Brucella broth. C) CryoEM images of *B. abortus* wild-type and Δ*eipB* cells cultivated in liquid broth that either remained untreated or were treated with 2 mM EDTA or 5 μg/ml ampicillin for 4 hours.

Although Δ*eipB* CFUs were reduced relative to wild-type on agar plates containing all three envelope stressors that we assayed, we observed no apparent defects in Δ*eipB* cell morphology by light microscopy or cryo-electron microscopy when cultivated in liquid broth (Figure 3B and C). Incubation of Δ*eipB* with 2 mM EDTA or 5 μg/ml ampicillin (final concentration) in Brucella broth for 4 hours also had no apparent effect on cell structure, nor did *eipB* overexpression (Figure 3B and C). Longer periods of growth in the presence of stressors may be required for differences in cell morphology/structure to be evident in broth. It may also be the case that the envelope stress phenotypes we observe are particular to growth on solid medium.

### *B. abortus* Δ*eipB* agglutination phenotypes indicate the presence of smooth LPS

In *B. abortus*, smooth LPS (containing O-polysaccharide) is an important virulence determinant (15). Smooth LPS can also act as a protective layer against treatments that compromise the integrity of the cell envelope (16). Loss of smooth LPS in *B. abortus* Δ*eipB* could therefore explain the phenotypes we observe for this strain. To test this hypothesis, we assayed wild-type and Δ*eipB* agglutination in the presence of serum from a *B. abortus*-infected mouse. A major serological response to smooth *Brucella* species is to O-polysaccharide (17), and thus agglutination can provide an indirect indication of the presence or absence of smooth LPS on the surface of the cell. Both wild-type and Δ*eipB* strains agglutinated in the presence of serum from a *B*. abortus-infected mouse, providing evidence for the presence of O-polysaccharide in Δ*eipB* (Figure S2A). As a negative control, we incubated the naturally rough species *B. ovis* with the same serum; *B. ovis* did not agglutinate in the presence of this serum (Figure S2A). We further assayed agglutination of *B. abortus* wild-type and Δ*eipB* strains in the presence of acriflavine, which is demonstrated to agglutinate rough strains such as *B. ovis* (18, 19). After 2 hours of incubation, we observed no agglutination of wild-type *B. abortus* or Δ*eipB* (Figure S2B). We treated *B. ovis* with acriflavine as a positive control and observed agglutination as expected (Figure S2B). Together, these data indicate that deletion of *eipB* does not result in a loss of smooth LPS. However, we cannot rule out the possibility that the chemical structure of O-polysaccharide is altered in Δ*eipB*.

### EipB is a monomeric protein that is secreted to the periplasm

The N-terminus (residues M1-A30) of *Brucella* EipB contains a predicted signal peptide based on SignalP 4.2 analysis (20). EipB (DUF1849) homologs in other *Alphaproteobacteria* also have a predicted N-terminal secretion signal (Figure S3). We note that EipB in our wild-type *B. abortus* 2308 strain has a methionine instead of a leucine at position 250. These two amino acids are interchangeable at this position in DUF1849 (Figure S4). To test the prediction that EipB is a periplasmic protein, we fused the *Escherichia coli* periplasmic alkaline phosphatase gene (*phoA*) to *B. abortus eipB* and expressed fusions from a *lac* promoter in *B. ovis*. We generated (i) the full-length EipB protein (M1-K280) fused at its C-terminus to *E. coli* PhoA (EipB-PhoA_Ec_) and (ii) an EipB-PhoA fusion lacking the hypothetical EipB signal peptide sequence (EipB^S29-K280^-PhoA_Ec_). After overnight growth in Brucella broth in presence or absence of 1 mM isopropyl **β**-D-1-thiogalactopyranoside (IPTG), we adjusted each culture to the same density and loaded into a 96-well plate containing 5-bromo-4-chloro-3-indolyl phosphate (BCIP, final concentration 200 μg/ml). BCIP is hydrolyzed to a blue pigment by PhoA, which can be measured colorimetrically. BCIP diffusion through the inner membrane is inefficient, and thus this reagent can be used to specifically detect PhoA activity in the periplasmic space or in the extracellular medium (21). After a 2-hour incubation at 37°C, the well containing the *B. ovis* cells expressing the EipB^M1-K280^-PhoA_Ec_ fusion turned dark blue. We observed no color change in the well containing the *B. ovis* strain expressing the EipB^S29-K280^-PhoA_Ec_ protein fusion (Figure 4A). As expected, no color change was observed in absence of induction with 1 mM IPTG (Figure 4A). To test if EipB is secreted from the cell into the growth medium, we performed a similar experiment on spent medium supernatants from the different cultures. We observed no color change in these samples after 2 hours of incubation providing evidence that EipB^M1-K280^-PhoA_Ec_ is not secreted from the cell.

**Figure 4:**
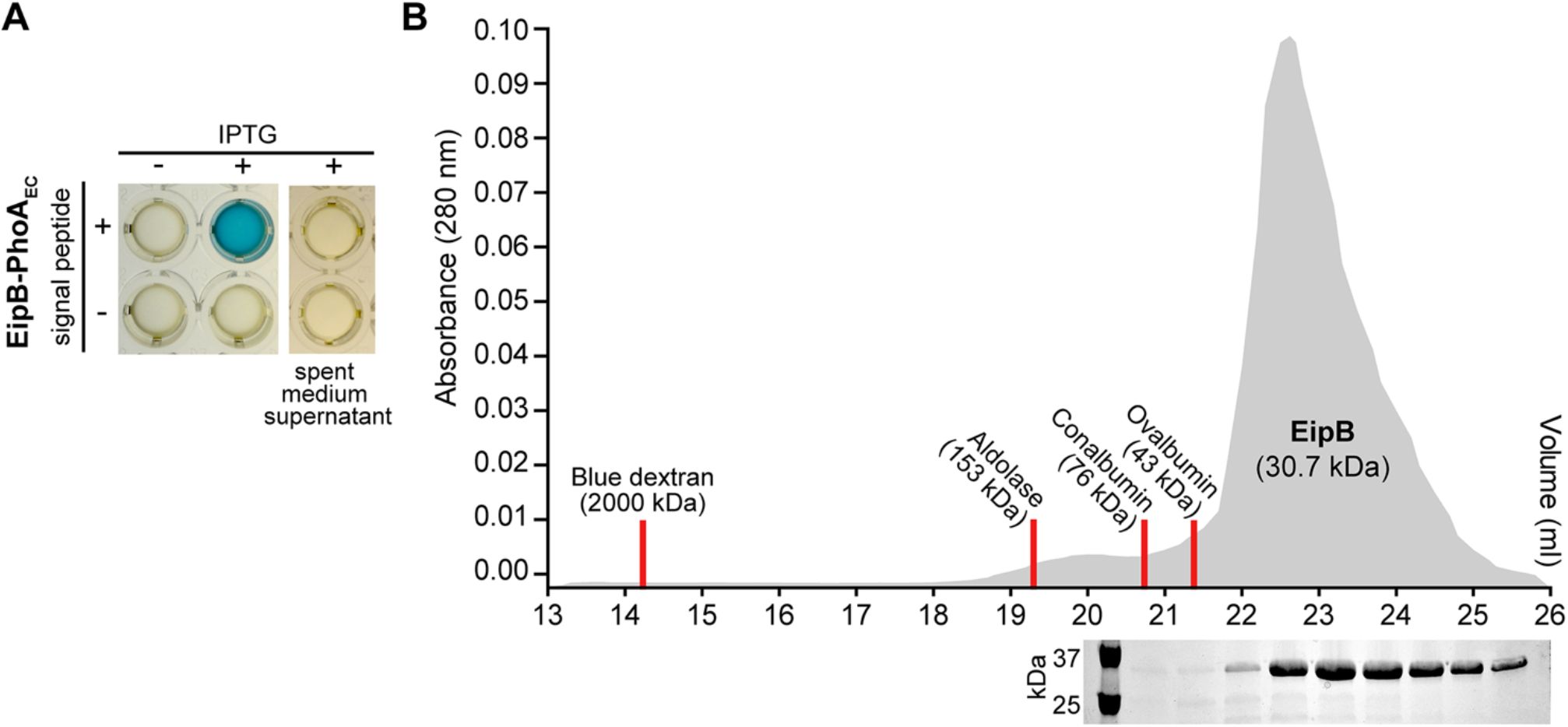
EipB is monomeric in solution and is secreted to the *Brucella* periplasm. A) Alkaline phosphatase assay. Overnight cultures of *B. ovis* expressing EipB with (+) or without (-) its signal peptide and fused to *E. coli* PhoA, were grown in presence (+) or absence (-) of 1 mM IPTG inducer. In a 96-well plate, these cultures were mixed with BCIP (200 μM final concentration) and developed for 2 hours at 37°C / 5% CO2. Only the strain expressing EipB-PhoA_Ec_ with a signal peptide turned blue, providing evidence that the protein is located in the periplasm. As a control, spent medium supernatants were mixed with BCIP to test whether EipB-PhoA_Ec_ is secreted into the medium. After 2 hours incubation, no color change was observed, indicating that EipB-PhoA_Ec_ is not exported outside the cell. These experiments were performed at least three times with independent clones. A representative image is shown. B) Size exclusion chromatography elution profile of purified EipB (in grey). Elution fractions were loaded on a SDS-PAGE, and a single band migrating at ~30 kDa was visible. Elution peaks of the molecular weight standards (blue dextran: 2000 kDa, aldolase: 157 kDa, conalbumin: 76 kDa, ovalbumin: 43 kDa) are shown as red line. This experiment was performed twice and yielded similar elution profiles.

We further assayed the oligomeric state of affinity-purified *B. abortus* EipB in solution by size-exclusion chromatography. The calculated molecular mass of His_6_-EipB (V31-K280) is 30.7 kDa. This protein eluted from a sizing column at a volume with an apparent molecular mass of ~23 kDa, which is consistent with a monomer (Figure 4B). There was no evidence of larger oligomers by size-exclusion chromatography. From these data, we conclude that EipB is a monomeric periplasmic protein.

### EipB folds into a spiral-like β-sheet that resembles PA1994, LolA and LolB

We postulated that the three-dimensional structure of EipB may provide molecular-level insight into its function in the cell. As such, we solved an x-ray crystal structure of *B. abortus* EipB (residues A30-K280; PDB ID: 6NTR). EipB lacking its signal peptide formed triclinic crystals (*a*=47.4 Å *b*=69.2 Å, *c*=83.2 Å, ⍰=90.1, ⍰=90.0°, ⍰=78.7°) that diffracted to 2.1 Å resolution; we refined this structure to Rwork= 0.195 and Rfree= 0.245. Crystallographic data and refinement statistics are summarized in Table S2. Four EipB molecules (chains A-D) are present in the crystallographic asymmetric unit.

Each EipB monomer consists of 14 antiparallel **β**-strands (**β**1-**β**14) forming an oval, spiral-like **β**-sheet (minor axis diameter: ~25 Å; major axis diameter: ~35 Å). Two regions of this **β**-spiral, involving **β**5, **β**6, **β**7, **β**8 and the hairpin loop connecting **β**9 and **β**10, overlap (Figure 5A and B). Interactions between these two overlapping portions of structure are mostly hydrophobic, though polar contacts are also found in these regions (Figures 5 and 6). One side of the spiral is occluded by the N-terminus, a loop connecting **β**-strands 12 and 13, and **α**-helix 1, which form the bottom of this “cup” shaped protein (Figures 5 and 6A). The external surface of EipB is positively and negatively charged, and also presents small hydrophobic patches (Figure S5); one helix, **α**2, is kinked and positioned at the surface of the cylindrical **β**-spiral (Figure 5A and B). The lumen of EipB is solvent accessible and is partially filled with the side chains of hydrophobic or acidic residues. Hydrophobic residues represent ~66% of the residues present inside the EipB cavity (Figures 5 and 6B). The size of this cavity suggests that EipB, in this conformation, can accommodate small molecules or ligands in its lumen.

**Figure 5:**
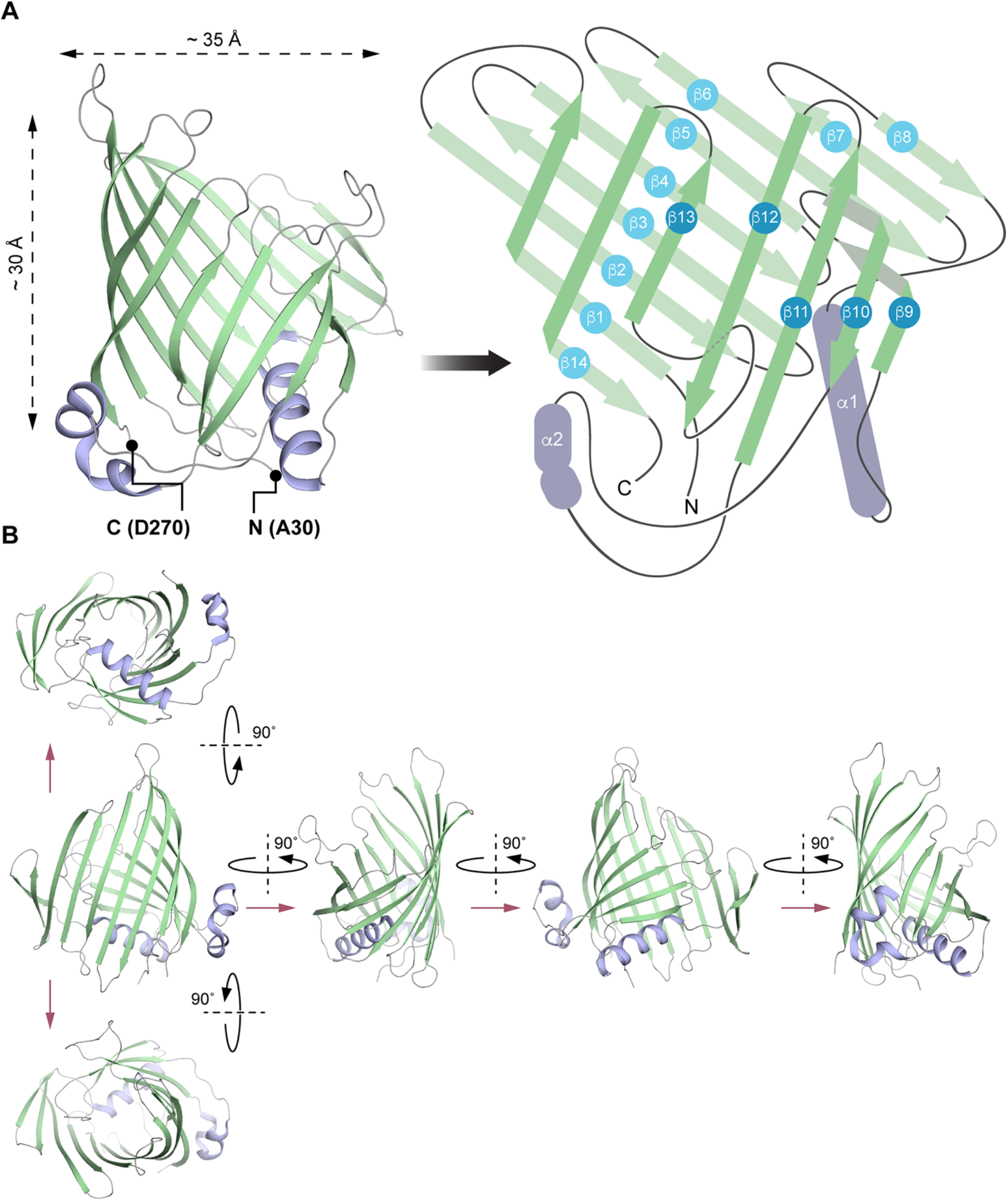
EipB adopts a **β**-spiral fold. A) Left: X-ray structure of EipB. EipB consist of 14 **β**-strands (in green) and 2 **α**-helices (in violet). The N-terminus (A30) and the C-terminus (D270) are reported on this structure. Right: simplified representation of EipB; color code is the same as before. B) Different orientations of EipB structure; color code is the same as before.

**Figure 6:**
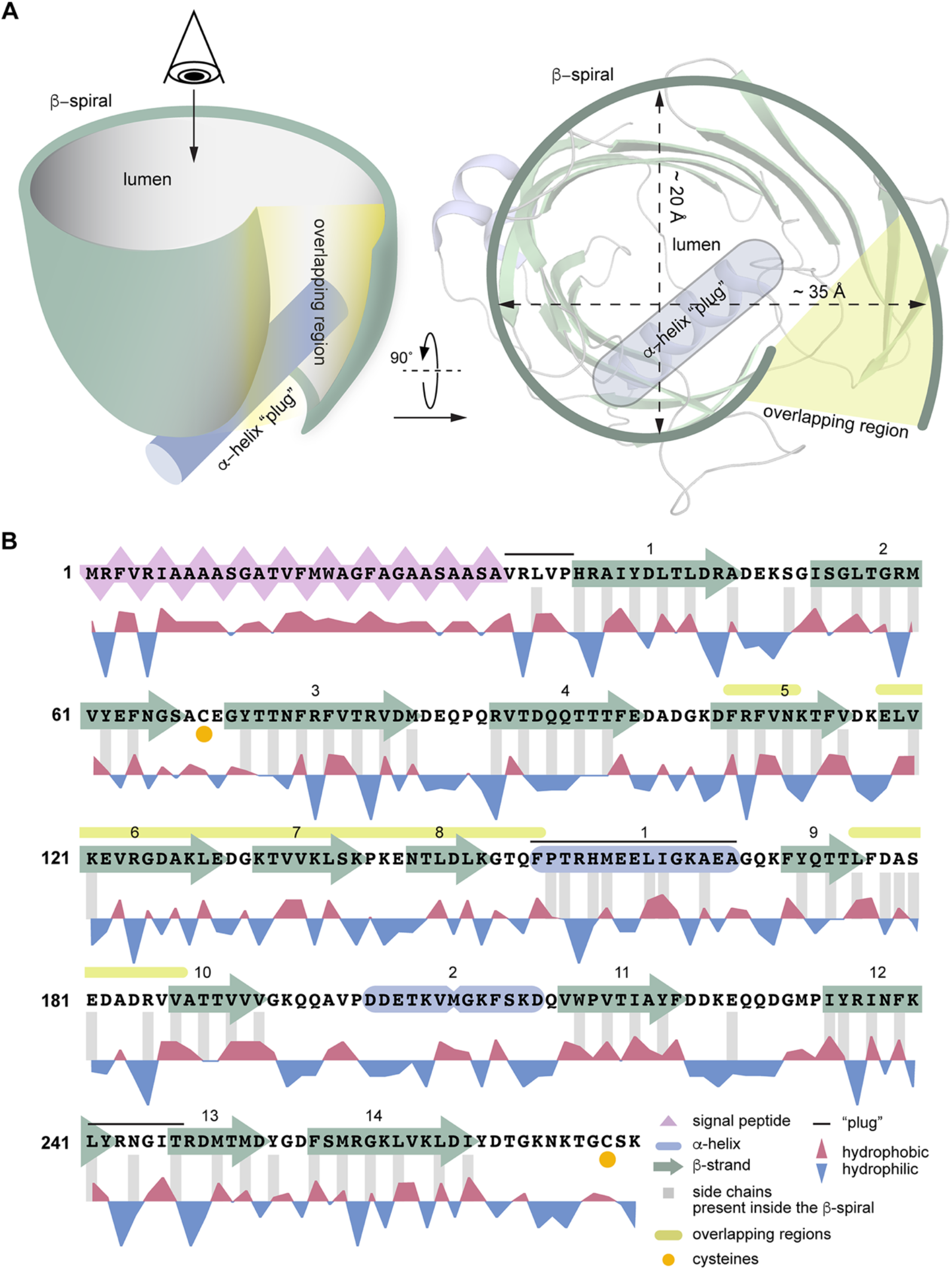
Simplified representation of EipB structure. A) EipB adopts a cup-like structure, fourteen **β**-strands (in green) form an overlapping **β**-spiral (**β**5-**β**6-**β**7-**β**8 overlap with **β**9-**β**10 connecting loop, highlighted in yellow in panel A and B). **α**1 (in violet) and the loop connecting **β**12 and **β**13 form the bottom of this “cup”. B) Amino acid sequence of EipB. The sequence corresponding to the predicted signal peptide is highlighted in pink. **β**-strands and **α**-helices are represented by green arrows and violet cylinders, respectively. Hydrophobic (red) and hydrophilic (blue) residues are reported below the sequence. Residues with side chains present inside EipB cavity are highlighted with grey bars. Cysteines C69 and C278 are highlighted with orange dots. Structural elements forming the bottom of the **β**-spiral are highlighted with a black line; overlapping regions are highlighted with a yellow line.

We searched the EipB structure against the protein structure database using Dali (22), but failed to identify clear structural homologs. *Pseudomonas aeruginosa* PA1994 (PDB ID: 2H1T) (23) was the closest structural match to EipB (RMSD ~3.5; Z-score ~11) (Figure S6A). Despite very low sequence identity (~8%), PA1994 has noticeable structural similarities to EipB: it adopts a spiral-like **β**-fold involving 15 **β**-strands, which is occluded at one end with a long **α**-helix. Unlike EipB, PA1994 lacks a signal peptide and is predicted to be a cytoplasmic protein. Structural parallels between PA1994 and the periplasmic lipoprotein chaperones LolA/LolB have been noted and a role for PA1994 in glycolipid metabolism has been postulated (23), though this prediction remains untested. Like PA1994, EipB has structural similarities to LolA and LolB, in particular the antiparallel and curved β-sheet scaffold that engulfs a central α-helical plug (Figure S6B). Whether *Brucella* EipB, or DUF1849 proteins more generally, function in trafficking lipoproteins or other molecules in the periplasm remains to be tested.

### EipB has a conserved disulfide bond

We identified two cysteines in EipB, C69 and C278, which are the two most conserved residues in the DUF1849 sequence family (Figures S3 and S4). C69 is solvent exposed in *Brucella* EipB and positioned in a loop connecting **β**2 and **β**3. C278 is present at the C-terminus of the protein, which immediately follows **β**14. **β**14 interacts with **β**13 and **β**1, and is spatially proximal to **β**2 and **β**3 (Figure 7A). Given the proximity of these two cysteines in the EipB structure, we hypothesized that C69 and C278 form an internal disulfide bond. However, electron density for the 10 C-terminal residues (containing C278) is not well resolved in the EipB crystal structure, and a disulfide bond is not evident, likely because the protein was dialyzed against a buffer containing 2 mM 1,4-dithiothreitol (DTT) prior to crystallization.

**Figure 7:**
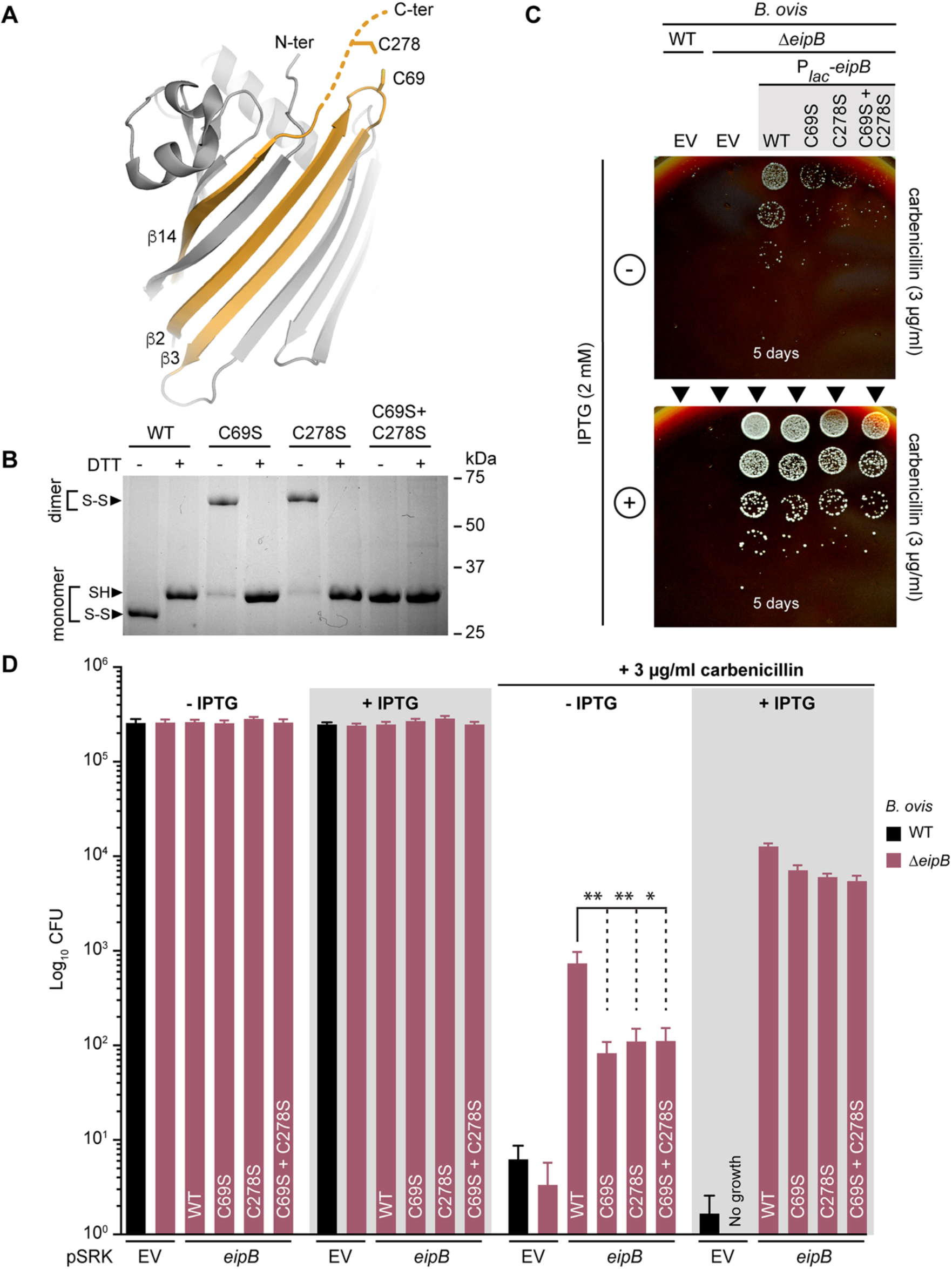
EipB has an internal disulfide bond. A) Cysteines C69 and C278 are spatially proximal in the EipB structure and form a disulfide bond. C278 is present at the EipB C-terminus that follows **β**14, and C69 is present in a loop connecting **β**2 and **β**3. B) His-tagged wild-type EipB and EipB cysteine mutant proteins (C69S, C278S, and C69S+C278S) were purified and mixed with a protein loading buffer plus or minus 1 mM DTT. Protein samples were resolved by 12% SDS-PAGE. This experiment was performed three times. Picture of a representative gel is presented. C) Growth on SBA plates containing 3 μg/ml of carbenicillin with (+) or without (-) 2 mM IPTG of a serially diluted (10-fold dilution) *B. ovis* Δ*eipB* strain ectopically expressing wild-type EipB (P_*lac*_-*eipB*), C69S mutant (P_*lac*_-*eipB*^C69S^), C278S mutant (P_*lac*_-*eipB*^C69S^), or C69S+C278S mutant (P*_lac_-eipB*^C69S+C278S^). *B. ovis* wild-type (WT) and Δ*eipB* carrying the pSRK empty vector (EV) were used as a control. Days of growth at 37°C / 5% CO_2_ are reported for each plate. A representative picture of the different plates is presented. D) Enumerated CFUs after growth on SBA plates containing 3 μg/ml of carbenicillin with (+) or without (-) 2 mM IPTG of serially diluted (10-fold dilution) *B. ovis* Δ*eipB* strains expressing different versions of *eipB* from a plasmid (wild-type and cysteine mutants; see panel C legend). Empty vector (EV) strains and SBA plates with no carbenicillin, plus or minus IPTG, were used as controls. This experiment was independently performed twice with two different clones each time, and all plate assays were done in triplicate. Each data point is the mean ± the standard error of the mean. One-way ANOVA followed by Dunnett’s post test (to wild-type) supports the conclusion that eipB-dependent protection against the cell wall antibiotic, carbenicillin, is significantly diminished when disulfide-forming residues C69 (**, p<0.005) and C278 (**, p<0.003) are individually or both (*, p<0.01) mutated to serine. This is effect is evident with leaky *eipB* expression from P_*lac*_, but diminished when expression of wild-type and mutant *eipB* alleles is induced by IPTG.

To biochemically test if these two cysteines form a disulfide bond, we purified *B. abortus* EipB under non-reducing conditions and mixed the protein with SDS gel loading dye with or without 1 mM dithithreitol (DTT). We observed two bands that migrated differently in the 30 kDa region when the protein was resolved by 12% SDS-PAGE. EipB without DTT migrated farther than the DTT-treated protein, suggesting the presence of a disulfide bond (Figure 7B). We performed this same experiment with three different EipB cysteine mutant proteins in which C69, C278, or both were mutated to serine. In the absence of DTT, EipB^C69S^ and EipB^C278S^ migrated at an apparent molecular weight of ~60 kDa, corresponding to a dimeric EipB interacting through a S-S bond. After DTT treatment, these mutant proteins migrated the same as the reduced wild-type protein (Figure 7B). As expected, the double cysteine mutant (EipB^C69S+C278S^) did not form an apparent dimer and was unaffected by DTT (Figure 7B). From these data, we conclude that an internal disulfide bond can form between C69 and C278 in EipB and is likely present *in vivo*, as EipB resides in the oxidizing environment of the periplasm.

To test whether this disulfide bond affects EipB function, we measured CFUs of a *Brucella ovis* Δ*eipB* (Δ*bov_1121*) strain expressing wild-type *B. abortus* EipB or cysteine disulfide mutants on agar plates containing 3 μg/ml carbenicillin. *B. ovis* is a closely related biosafety level 2 (BSL2) surrogate for *B. abortus. B. ovis* and *B. abortus* EipB are identical with the exception of one amino acid at position 250 (Figure S4). In this carbenicillin assay (Figure 7C and D), *B. abortus* EipB complemented a *B. ovis* Δ*eipB* strain, suggesting that the substitution at residue 250 does not impair EipB function. We placed four different versions of *eipB* under the control of a *lac* promoter (P_*lac*_): P_*lac*_-*eipB*^WT^, P_*lac*_-*eipB*^C69S^, P_*lac*_-*eipB*^C278S^, and P_*lac*_-*eipB*^C69S+C278S^; the empty vector was used as a control. After 5 to 6 days of growth on Schaedler Blood Agar (SBA) plates containing 3 μg/ml of carbenicillin and no IPTG, we observed poor growth at only the lowest dilution for wild-type and Δ*eipB* strains carrying the empty vector control (also see Figure S7A for an example of growth on 2 μg/ml carbenicillin plates). Corresponding colonies for the strains carrying the different P_*lac*_-*eipB* overexpression plasmids were more abundant though very small in the absence of IPTG induction. However, the strain harboring the wild-type *eipB* plasmid systematically grew at 1 log higher dilution than the cysteine mutant strains indicating that the presence of the disulfide bond in *eipB* contributes to carbenicillin resistance on solid medium (Figure 7C and D, see also Figure S7A). These results indicate some level of leaky expression from the multi-copy P_*lac*_-*eipB* plasmids. When induced with IPTG, overexpression of the different EipB variants enhanced growth in all strains. (Figure 7C and D). As expected, strains grown on control plates without carbenicillin had no growth defect, with or without IPTG induction (Figure 7D). The morphology of *B. ovis* Δ*eipB* strains expressing the different variants of *eipB* appeared normal by phase contrast microscopy (see Figure S7B). These results provide evidence that EipB is necessary for full carbenicillin resistance in *B. ovis*, and that cysteines 69 and 278 contribute to EipB function *in vivo*.

To evaluate the effect of these two cysteines on EipB stability *in vitro*, we measured the thermal stability of purified wild-type *B. abortus* EipB (EipB^WT^) and double cysteine mutant (EipB^C69S+C278S^) in presence or absence of 2 mM DTT. EipB^WT^ melted at ~46°C in absence of DTT and at ~41.5°C in presence of DTT. EipB^C69S+C278S^ melted at ~42.3°C in the presence or absence of DTT (see Figure S8). We conclude that an internal disulfide bond stabilizes EipB structure *in vitro*. Reduced stability of EipB lacking its conserved disulfide bond may contribute to the 1 log relative growth defect of Δ*eipB* strains expressing EipB cysteine mutants on SBA carbenicillin plates (Figure 7C and D).

### *eipB* deletion is synthetically lethal with *bab1_0430* (*ttpA*) disruption, and synthetically sick with disruption of multiple genes with cell envelope functions

To further characterize how *eipB* functions in the *Brucella* cell, we aimed to identify transposon (Tn) insertion mutations that are synthetically lethal with *eipB* deletion in *B. abortus* (see Tables S3 and S4). In other words, we sought to discover genes that are dispensable in a wild-type genetic background, but that cannot be disrupted in a Δ*eipB* background. By sequencing a Tn-Himar insertion library generated in *B. abortus* Δ*eipB* (NCBI Sequence Read Archive accession SRR8322167) and a Tn-Himar library generated in wild-type *B. abortus* (NCBI Sequence Read Archive accession SRR7943723), we uncovered evidence that disruption of *bab1_0430* (RefSeq locus BAB_RS17965) is synthetically lethal with *eipB* deletion. Specifically, reproducible reads corresponding to insertions in the central 10-90% of *bab1_0430* were not evident in Δ*eipB*, but were present in wild-type (Figure 8A). *bab1_0430* encodes a 621-residue **<u>t</u>**etratrico**<u>p</u>**eptide **<u>r</u>**epeat-containing (TPR) protein with a predicted signal peptide and signal peptidase site at its N-terminus. This protein was previously detected by mass spectrometry analyses of *B. abortus* extracts, and described as a cell-envelope associated (24), or periplasmic protein (25). Hereafter, we refer to this gene as *ttpA* (**<u>t</u>**etra**<u>t</u>**ricopeptide repeat **<u>p</u>**rotein **<u>A</u>**) based on its similarity to *Rhizobium leguminosarum ttpA* (12).

**Figure 8:**
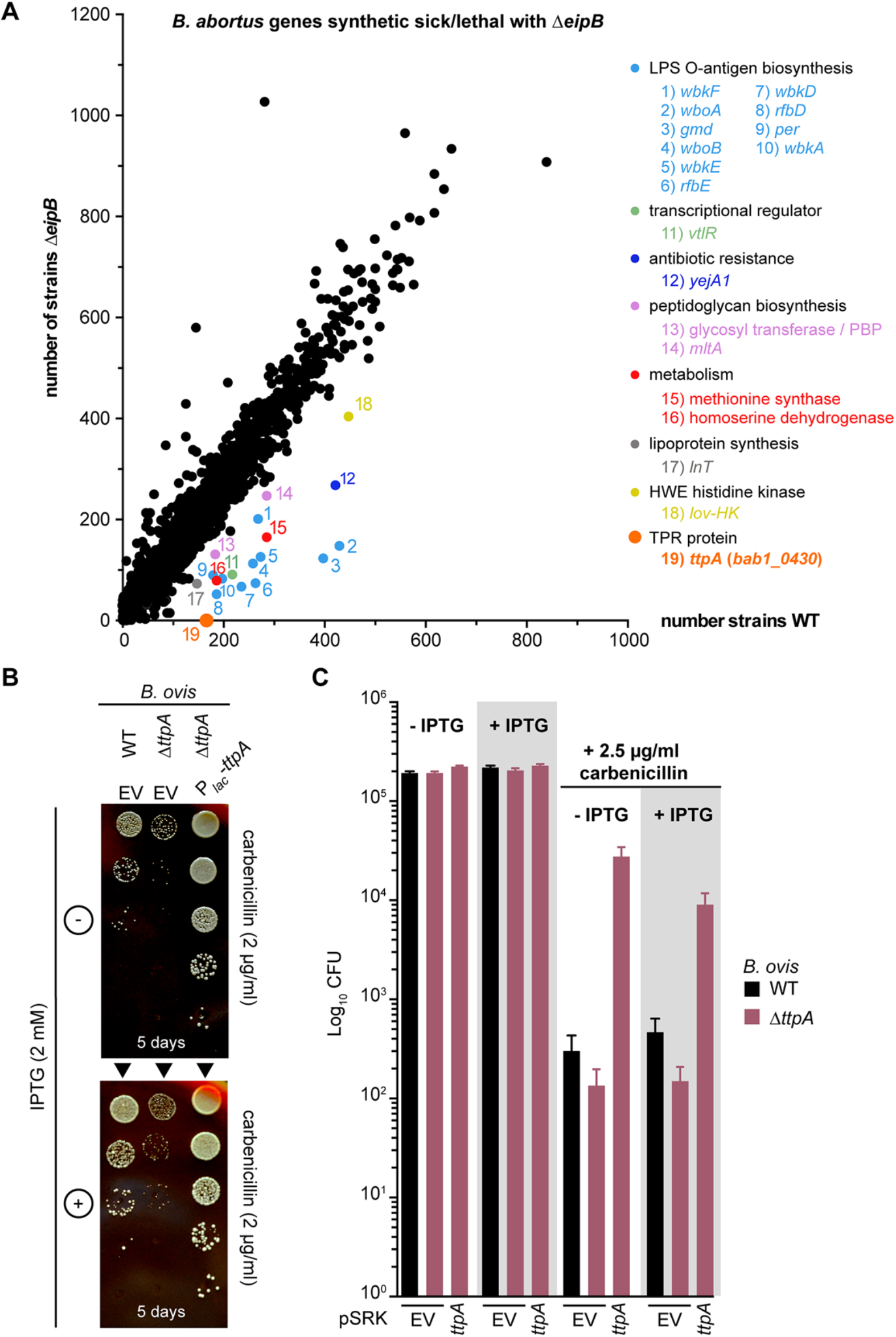
*B. abortus eipB* deletion is synthetically lethal with Tn-Himar disruption of *bab1_0430*, which encodes a tetratricopeptide repeat (TPR) protein. A) Identification of *B. abortus* genes that are synthetically lethal or sick with *eipB* deletion. Tn-Himar insertion strains per gene (black dots) obtained in a *B. abortus* Δ*eipB* background are plotted as a function of strains per gene in a wild-type background. *bab1_0430*, for which we observed significantly fewer insertions in Δ*eipB* than in wild-type, is represented as an orange dot. Other synthetic sick genes are also evident in the plot, including genes involved in LPS O-antigen synthesis in light-blue: *wbkF* (locus *bab1_0535*); *wboA* (*bab1_0999*); *gmd* (*bab1_0545*); *wboB* (*bab1_1000*); *wbkE* (*bab1_0563*); *rfbE* (*bab1_0542*); *wbkD* (*bab1_0534*); *rfbD* (*bab1_0543*); *per* (*bab1_0544*); *wbkA* (*bab1_0553*). Genes related to peptidoglycan synthesis in pink: *mltA* (*bab1_2076*); penicillin-binding protein (*bab1_607*). Apolipoprotein N-acyltransferase *lnt* (*bab1_2158*) is in grey; LysR transcriptional regulator *vtlR* (*bab1_1517*) is in light green; extracellular solute binding protein *yejA1* (*bab1_0010*) is in dark blue; general stress response kinase *lovhK* (*bab2_0652*) is in yellow; metabolic genes methionine synthase (*bab1_0188*) and homoserine dehydrogenase (*bab1_1293*) are in red. B) Growth on SBA plates containing 2 μg/ml of carbenicillin ± 2 mM IPTG of serially diluted (10-fold dilution) *B. ovis* Δ*ttpA* strains carrying the pSRK empty vector (EV) or ectopically expressing wild-type TtpA (P_*lac*_-*ttpA*). The wild-type (WT) *B. ovis* pSRK empty vector (EV) strain was used as a control. Days of growth at 37°C / 5% CO2 are reported for each plate. A representative picture of the different plates is presented. C) Enumerated CFUs, after growth on SBA plates containing 2.5 μg/ml of carbenicillin ± 2 mM IPTG, of serially diluted (10-fold dilution) *B. ovis* wild-type (black) and Δ*ttpA* (dark pink) strains. The Δ*ttpA* strain was either transformed with the empty vector (EV) or with pSRK-ttpA. Empty vector (EV) wild-type strain and SBA plates with no carbenicillin, and plus or minus IPTG were used as controls. This experiment was independently performed twice with two different clones each time, and all plate assays were done in triplicate. Each data point is the mean ± the standard error of the mean.

Genes involved in LPS O-antigen synthesis, and previously described as synthetic lethal with *eipA* (*bab1_1612*) deletion in *B. abortus* (8), were synthetic sick with *eipB* deletion (Figure 8A), as were genes involved in peptidoglycan synthesis: *mltA (bab1_2076*, lytic murein transglycosylase A) and *bab1_0607* (glycosyl transferase/penicillin-binding protein 1A) (26) (Figure 8A). There were reduced transposon insertions in solute binding protein *yejA1* (*bab1_0010*) (Figure 8A), which is involved in *B. melitensis* resistance to polymyxin (27). *lnt* (*bab1_2158*) and *vtlR* (*bab1_1517*) were also synthetic sick with Δ*eipB. lnt* is an apolipoprotein N-acyltransferase involved in lipoprotein synthesis (28); *vtlR* encodes a LysR transcriptional regulator required for full *B. abortus* virulence (29) (Figure 8A). Finally, the general stress sensor kinase *lovHK* (*bab2_0652*) (30), *bab1_1293* (homoserine dehydrogenase), and *bab1_0188* (methionine synthase), had fewer Tn insertions in the Δ*eipB* background relative to wild-type (Figure 8A).

### ttpA contributes to carbenicillin resistance

As *ttpA* disruption is synthetic lethal with *eipB* deletion, we postulated that these two genes have complementary functions or are involved in a common physiological process (i.e. envelope integrity). Thus, to characterize *ttpA* and the nature of its connection to *eipB*, we deleted *ttpA* in *B. ovis* and evaluated its sensitivity to carbenicillin. All efforts to delete *B. ovis ttpA* (locus tag *bov_0411*) using a classic crossover recombination and *sacB* counterselection approach were unsuccessful, though hundreds of clones were screened. Efforts to delete the chromosomal copy by expressing a copy of *ttpA* from a plasmid also failed. This result is surprising considering that transposon insertions in *B. abortus ttpA* (NCBI Sequence Read Archive accession SRR7943723) and *B. ovis ttpA* (NCBI Sequence Read Archive accession SRR7943724) are tolerated in wild-type backgrounds (8). As an alternative approach to study the function of this gene, we inactivated *ttpA* using a single crossover recombination strategy. The resulting strain expressed a truncated version of TtpA containing the first 205 amino acids (including the signal peptide), immediately followed by 22 amino acids form the suicide plasmid. The corresponding *B. ovis* strain (Δ*ttpA*) was then transformed with a plasmid-borne IPTG-inducible copy of *ttpA* (pSRK-ttpA) or with an empty plasmid vector (EV). We evaluated sensitivity of these strains to carbenicillin by plating a dilution series on SBA plates containing 2 or 2.5 μg/ml carbenicillin, with or without IPTG inducer (Figure 8B and C). When compared to wild-type with empty vector, *B. ovis* Δ*ttpA* with empty vector had ~0.5 log reduced CFUs on carbenicillin SBA. The corresponding colonies of *B. ovis ΔttpA* were noticeably smaller than wild-type. Genetic complementation of Δ*ttpA* with pSRK-*ttpA* restored growth on carbenicillin plates. *B. ovis* Δ*ttpA*/pSRK-*ttpA* had ~1.5 log more colonies than wild-type in the presence of carbenicillin, with or without IPTG induction. Thus, leaky expression of *ttpA* from the *lac* promoter on pSRK-*ttpA* is apparently sufficient to protect this strain from carbenicillin on solid medium. Morphology of the *B. ovis* Δ*ttpA* strains appeared normal by phase contrast microscopy at 630x magnification (Figure S9).

To further evaluate the effect of *ttpA* overexpression, we assayed *B. ovis* wild-type and Δ*eipB* strains carrying pSRK-ttpA. As before, we tested sensitivity of these inducible expression strains to carbenicillin by plating a dilution series on SBA plates containing 3 μg/ml of carbenicillin, with or without 2 mM IPTG inducer (Figure 9A and B). Wild-type *B. ovis*/pSRK-*ttpA* and wild-type *B. ovis*/pSRK-*eipB* strains had equivalent CFUs in the absence of carbenicillin, with or without IPTG. *ttpA* or *eipB* provided a ~3 log protective effect without IPTG induction in the presence of carbenicillin compared to the wild-type empty vector strain (Figure 9). Surprisingly, inducing *ttpA* expression with IPTG reduced its ability to protect in the presence of carbenicillin by 1 log (relative to uninduced), and the corresponding colonies were very small suggesting slower growth when *ttpA* was induced (Figure 9A and B). This may be an effect of IPTG, based on reduced CFU counts of wild-type empty vector control under this condition. As expected, induced expression of *eipB* from P_*lac*_-*eipB* rescued the carbenicillin viability defect of Δ*eipB*. However, induced expression of *ttpA* from P_*lac*_-*ttpA* was not sufficient to rescue the Δ*eipB* carbenicillin phenotype (Figure 9A and B). As before, we observed highly reduced CFUs for *B. ovis* wild-type or Δ*eipB* control strains carrying the pSRK empty vector (EV), when challenged with 3 μg/ml of carbenicillin. Morphology of wild-type or Δ*eipB B. ovis* strains overexpressing *ttpA* appeared normal by phase contrast microscopy at 630x magnification (Figure S10).

**Figure 9:**
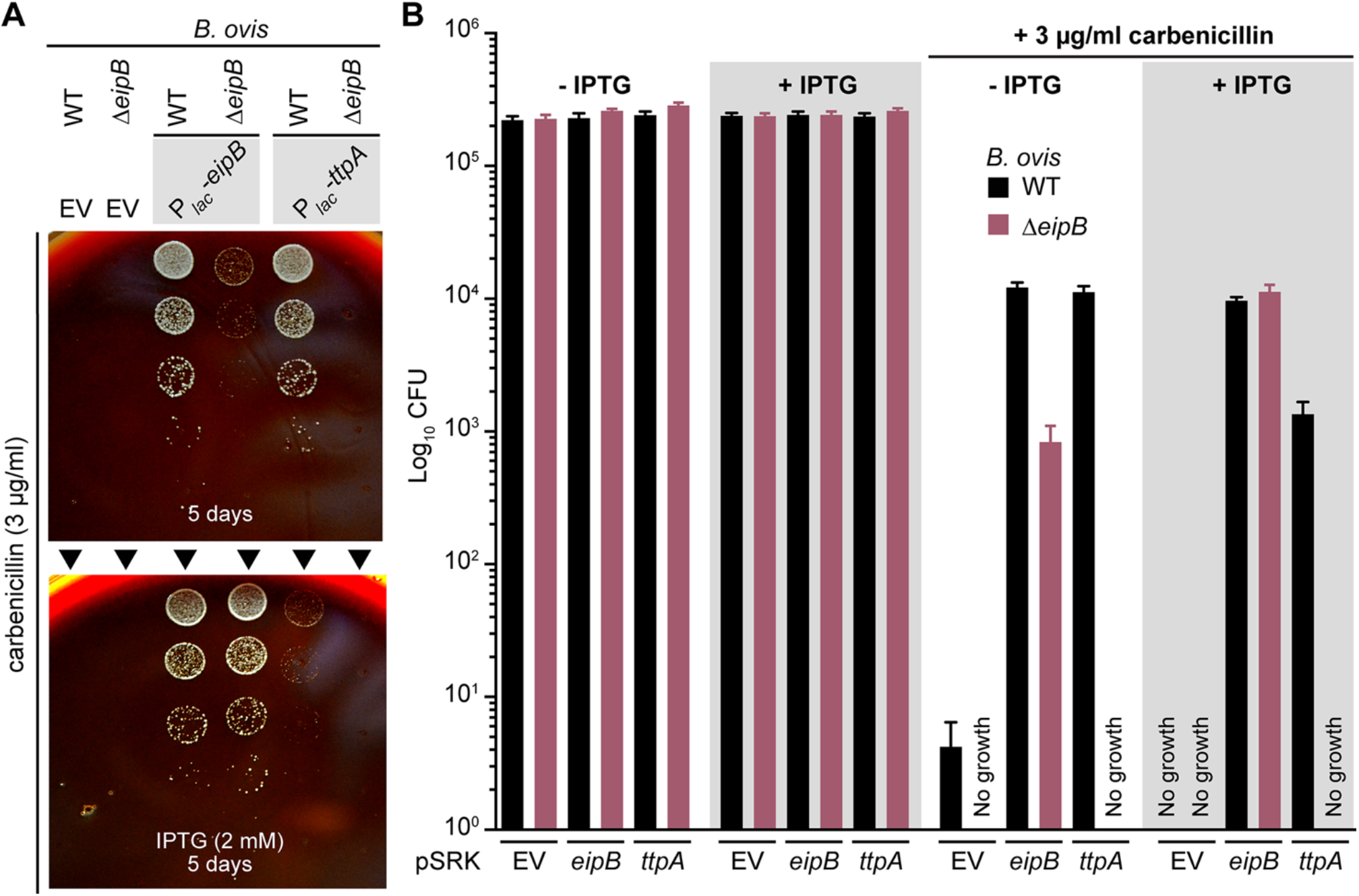
Overexpression of TtpA protects against carbenicillin treatment; protection requires EipB. A) Growth on SBA plates containing 3 μg/ml of carbenicillin ± 2 mM IPTG of serially diluted (10-fold dilution) *B. ovis* wild-type (WT) and Δ*eipB* strains expressing wild-type EipB (P_*lac*_-*eipB*) or TtpA (P_*lac*_-*ttpA*). *B. ovis* strains carrying the pSRK empty vector (EV) were used as a control. Days of growth at 37°C / 5% CO2 are reported for each plate. A representative picture of the different plates is presented. B) Enumerated CFUs after growth on SBA plates containing 3 μg/ml of carbenicillin ± 2 mM IPTG of serially diluted (10-fold dilution) *B. ovis* wild-type (black) and Δ*eipB* (dark pink) strains ectopically expressing *eipB* or *ttpA*. Empty vector (EV) strains and SBA plates with no carbenicillin, and plus or minus IPTG were used as controls. This experiment was independently performed twice with two different clones each time, and all plate assays were done in triplicate. Each data point is the mean ± the standard error of the mean.

The observed genetic interaction between *eipB* and *ttpA*, the fact that both single mutants have envelope phenotypes, and the fact that both gene products are secreted to the periplasm raised the possibility that EipB and TtpA physically interact. We tested interaction between EipB and TtpA proteins using bacterial two-hybrid and biochemical pull-down assays. We further evaluated whether a possible EipB-TtpA interaction is influenced by the presence or absence of the EipB internal disulfide bond using a biochemical pull-down. For our bacterial two-hybrid assay, EipB^V31-K280^ was fused to the T25 adenylate cyclase fragment, and TtpA^K31-D621^ was fused to the T18 or T18C adenylate cyclase fragments. For the pull-down assay, MBP-tagged TtpA (K31-D621) and His-tagged EipB (V31-K280; wild-type and the different cysteine mutants) were co-purified in presence or absence of DTT. We found no evidence for direct interaction between EipB and TtpA, suggesting that the function of these two proteins in *Brucella* envelope stress adaptation is not achieved through direct interaction (Figure S11).

## DISCUSSION

Bacterial genome sequencing efforts over the past two decades have revealed thousands of protein domains of unknown function (DUFs). The DUF1849 sequence family is prevalent in orders *Rhizobiales, Rhodobacterales* and *Rhodospirillales*. To date, the function of DUF1849 has remained undefined. We have shown that a DUF1849 gene in *Brucella* spp., which we have named *eipB*, encodes a 14-stranded **β**-spiral protein that is secreted to the periplasm. *eipB* is required for maintenance of *B. abortus* spleen colonization in a mouse model of infection (Figure 2), and *eipB* deletion in *B. abortus* and in *B. ovis* results in sensitivity to treatments that compromise the integrity of the cell envelope *in vitro* (Figure 3). Envelope stress sensitivity of the *B. abortus* Δ*eipB* mutant likely contributes to its reduced virulence in a mouse. We further demonstrate that EipB contains a conserved disulfide bond that contributes to protein stability and function *in vitro;* the importance of this conserved disulfide to EipB function *in vivo* remains to be determined (Figures 6, 7, S3 and S4)

### A lipoprotein connection?

An x-ray crystal structure of EipB shows that this periplasmic protein adopts an unusual **β**-spiral fold that shares structural similarity (DALI Z-score= 11.0) with a functionally-uncharacterized *P. aeruginosa* protein, PA1994, despite low sequence identity (Figure S6). It was previously noted (23) that PA1994 has structural features that resemble the lipoprotein carrier and chaperone proteins LolA and LolB, which have a central role in lipoprotein localization in select Gram-negative bacteria (31). Like LolA, LolB, and PA1994, *Brucella* EipB forms a curved hydrophobic **β**-sheet that is wrapped around an **α**-helix (Figure S6B). Homologs of LolA are present in *Brucella* and other *Alphaproteobacteria*, but homologs of LolB are missing (28). Given the EipB structure, its periplasmic localization, and the phenotypes of a Δ*eipB* deletion strain, it is tempting to speculate that EipB (DUF1849) has a LolB-like function in the *Brucella* cell. However, it seems unlikely that LolB and EipB function in a structurally- or biochemically-equivalent manner. Certainly, we observe surface-level similarity between LolA/LolB and EipB structures (Figure S6), particularly in the antiparallel β-sheet region, but these proteins have topological differences that distinguish their folds. Moreover, LolB is a membrane anchored lipoprotein that facilitates lipoprotein targeting at the inner leaflet of the outer membrane. In contrast, *Brucella* EipB does not have a predicted site for lipidation (i.e. a lipobox), and is therefore unlikely to function as a membrane-anchored protein. The number of unique barcoded Tn-Himar insertions in the apolipoprotein N-acyltransferase *lnt* (*bab1_2158; lnt* conserved domain database score < e^−173^) is lower than expected in a Δ*eipB* background relative to wild-type (Figure 8A). This provides indirect evidence for a link between *eipB* and lipoproteins. Lnt catalyzes the final acylation step in lipoprotein biogenesis (32), which is often considered to be an essential cellular process. However, like *Francisella tularensis* and *Neisseria gonorrhoeae* (33), *B. abortus lnt* is dispensable (26) (Figure 8A and Table S4). The data presented here suggest that transposon insertions are less tolerated in *B. abortus lnt* when *eipB* is missing. Additional experimentation is required to test a possible functional relationship between *lnt* and *eipB*. However, it is notable that we did not observe a synthetic genetic interaction between *lnt* and the gene encoding a structurally-unrelated periplasmic envelope integrity protein, EipA, in a parallel Tn-seq experiment (8). Whether *eipB* actually influences lipoprotein biogenesis or localization remains to be tested.

### TtpA: a periplasmic determinant of cell envelope function in Rhizobiaceae

Transposon disruption of *ttpA* (*bab1_0430*) is not tolerated when *eipB* is deleted in *B. abortus. ttpA*, like *eipB*, contributes to carbenicillin resistance *in vitro* (Figures 8 and 9). Though we observed a genetic interaction between *eipB* and *ttpA*, we found no evidence for a direct physical interaction between the two periplasmic proteins encoded by these genes (Figure S11). TtpA is named for its tetratricopeptide repeat (TPR) motif; proteins containing TPR motifs are known to function in many different pathways in bacteria including cell envelope biogenesis, and are often molecular determinants of virulence (34, 35). Indeed, deletion of *ttpA* has been reported to attenuate *B. melitentis* virulence in a mouse infection model of infection (11) and to increase *R. leguminosarum* membrane permeability and sensitivity to SDS and hydrophobic antibiotics (12). A genetic interaction between *ttpA* and the complex media growth deficient (*cmdA-cmdD*) operon has been reported in *R. leguminosarum*. Mutations in this operon result in envelope dysfunction and defects in cell morphology (12, 36). While *B. abortus* contains a predicted *cmd* operon (*bab1_1573, bab1_1574, bab1_1575, and bab1_1576*) these genes remain uncharacterized. We found no evidence for a synthetic genetic interaction between *eipB* and *cmd* in *B. abortus*.

Leaky expression of either *eipB* or *ttpA* from a plasmid strongly protected *B. ovis* from a cell wall antibiotic (carbenicillin). Surprisingly, inducing *ttpA* expression from a plasmid with IPTG did not protect as well as uninduced (i.e. leaky) *ttpA* expression (Figure 9A and B). IPTG induction of *eipB* expression from a plasmid did not have this same parabolic effect on cell growth/survival in the face of carbenicillin treatment. Considering that EipB and TtpA confer resistance to **β**-lactam antibiotics, which perturb peptidoglycan synthesis, one might hypothesize that these proteins influence the structure or synthesis of the cell wall. This hypothesis is reinforced by the fact that a lytic murein transglycosylase and a class A PBP/glycosyl transferase are synthetic sick with *eipB* deletion (Figure 8A). In *E. coli*, the TPR-containing protein LpoA is proposed to reach from the outer membrane through the periplasm to interact with the peptidoglycan synthase PBP1A (37). Models in which EipB and TtpA influence lipoprotein biosynthesis and/or cell wall metabolism are important to test as we work toward understanding the mechanisms by which these genes ensure *Brucella* cell envelope integrity and survival in a mammalian host.

## Materials and Methods

Agglutination assays, mouse and macrophage infection assays, antibody measurements, and the transposon sequencing experiments for this study were performed in parallel with our recent studies of *eipA* (8).

All experiments using live *B. abortus* 2308 were performed in Biosafety Level 3 facilities according to United States Centers for Disease Control (CDC) select agent regulations at the University of Chicago Howard Taylor Ricketts Laboratory. All the *B. abortus* and *B. ovis* strains were cultivated at 37°C with 5% CO2; primer and strain information are available in Table S5.

### Chromosomal deletions in *B. abortus* and in *B. ovis*

The *B. abortus and B. ovis* Δ*eipB* deletion strains were generated using a double crossover recombination strategy as previously described (8). Briefly, fragments corresponding to the 500-base pair region upstream of the *eipB* start codon and the 500-base pair region downstream of the *eipB* stop codon were ligated into the suicide plasmid pNPTS138, which carries the *nptI* gene for initial kanamycin selection and the *sacB* gene for counter-selection on sucrose. Genetic complementation of the *B. abortus* deletion strain was carried out by transforming this strain with a pNPTS138 plasmid carrying the wild-type allele. The *B. ovis* Δ*eipB* strain was complemented with the pSRK-eipB plasmid (IPTG inducible).

To inactivate *ttpA* in *B. ovis* (*bov_0411*), a 527-nucleotide long internal fragment was cloned into pNPTS138-cam (a suicide plasmid that we engineered to carry a chloramphenicol resistance marker) and used to disrupt the target gene by single crossover insertion. The recombinant clones were selected on SBA plates supplemented with 3 μg/ml chloramphenicol. The corresponding strain expresses the first 205 amino acids (including the signal peptide) of TtpA, plus 22 extra amino acids from the plasmid sequence, followed by a stop codon. This Δ*ttpA* strain was complemented with pSRK-ttpA (kanamycin resistant).

### *Brucella* EipB and TtpA overexpression strains

For ectopic expression of *B. ovis* TtpA and the different versions of *B. abortus* EipB (wild-type, cysteine mutants, and the EipB-PhoA_Ec_ fusion with or without the signal peptide), the pSRKKm (Kan^R^) IPTG inducible plasmid was used (38). An overlapping PCR strategy was used to introduce cysteine mutations and to stitch the different DNA fragments to the *E. coli* alkaline phosphatase *phoA* (lacking its signal peptide). A Gibson-assembly cloning strategy was then used to insert the different DNA fragments in the linearized pSRK plasmid. After sequencing, plasmids were introduced in *B. abortus* or *B. ovis* by overnight mating with *E. coli* WM3064 in presence of 300 μM of diaminopimelic acid (DAP) and plated on SBA plates supplemented with kanamycin.

### Building and mapping the wild-type *B. abortus* and *B. abortus* Δ*eipB* Tn-Himar insertion libraries

To build and map the different Tn-Himar insertion libraries, we used a barcoded transposon mutagenesis strategy developed by Wetmore and colleagues (39). A full and detailed protocol can be found in our previous paper (8). Statistics for the two different transposon insertion libraries are reported in Table S3. For each Himar insertion library, Tn-seq read data have been deposited in the NCBI sequence read archive: *B. abortus* 2308 wild-type (BioProject PRJNA493942; SRR7943723), *B. abortus* Δ*eipB* (Δ*bab1_1186*) (BioProject PRJNA510139; SRR8322167).

### Cell culture and macrophage infection assays

Infection of inactivated macrophages differentiated from human monocytic THP-1 cells were performed as previously described (8). Briefly, for infection assays, 5 x 10^6^ *B. abortus* cells were used to infect 5 x 10^4^ THP-1 cells (multiplicity of infection of 1:100). To determine the numbers of intracellular bacteria at 1, 24 and 48 hours post-infection, the infected cells were lysed, the lysate was then serially diluted (10-fold serial dilution) and plated on TSA plates to enumerate CFUs.

### Mouse infection assay

All mouse studies were approved by the University of Chicago Institutional Animal Care and Use Committee (IACUC) and were performed as previously published (8). Briefly, 100 μl of a 5 x 10^5^ CFU/ml *B. abortus* suspension were intraperitoneally injected into 6-week-old female BALB/c mice (Harlan Laboratories, Inc.). At 1, 4, and 8 weeks post-infection, 5 mice per strain were sacrificed, and spleens were removed for weighing and CFU counting. At week 8, blood was also collected by cardiac-puncture and serum from each mouse was separated from blood using a serum separation tube (Sarstedt). Sera were subsequently used for Enzyme-Linked ImmunoSorbent Assays (ELISA).

### Determination of antibody responses at 8 weeks post infection

Total mouse serum IgG, IgG1, and IgG2a titers were measured using mouse-specific ELISA kits by following manufacturer’s instructions (eBioscience). *Brucella*-specific IgG titers were determined as previously published (8).

### Spleen histology

At 8 weeks post infection, spleens (n= 1 per strain) were prepared for histology as previously described (8). Briefly, spleens were first fixed with formalin and submitted for tissue embedding, Hematoxylin and Eosin (H & E) staining, and immunohistochemistry to Nationwide Histology (Veradale, Washington). For immunohistochemistry, goat anti-*Brucella* IgG was used (Tetracore, Inc). Pictures of fixed mouse spleen slides were subsequently analyzed and scored.

### Plate stress assays

Stress assays were performed as previously published (8). Briefly, the different *B. abortus* and *B. ovis* strains were resuspended in sterile PBS or Brucella broth to an OD600 of ~ 0.015 (~ 1 x 10^8^ CFU/ml) and serially diluted (10-fold serial dilution). 5 μl of each dilution were then spotted on TSA or SBA plates containing the different membrane stressors (2 to 5 μg/ml of ampicillin or carbenicillin, 200 μg/ml of deoxycholate or 2 mM EDTA final concentration).

To grow *B. ovis* strains containing pSRK-derived plasmids, all liquid cultures and plates were supplemented with 50 μg/ml kanamycin. When necessary, 2 mM IPTG (final concentration) was added to the plates to induce expression of EipB or TtpA from pSRK. We note that the *B. ovis* Δ*ttpA* strains carry the pNPTS138 suicide plasmid (used for gene disruption) which results in chloramphenicol resistance. However, no chloramphenicol was added to the overnight cultures or the stress plates. For carbenicillin growth/survival assays, *B. ovis* strains were grown for 3 days at 37°C / 5% CO2 on SBA plates without carbenicillin, and for 5 to 6 days when these plates contained 2, 2.5 or 3 μg/ml of carbenicillin.

### Cryo-electron microscopy

Cryo-electron microscopy was performed as previously described (8). Briefly, *B. abortus* cultures in Brucella broth (OD_600_ of ~0.015) were prepared with 2 mM EDTA or ampicillin (5 μg/ml) (final concentrations). After 4 hours of incubation in the presence of EDTA or ampicillin, cells were harvested and fixed in PBS + 4% formaldehyde. After 1 hour, cells were pelleted and resuspended in 500 μl EM buffer (40). Per CDC guidelines, cell killing was confirmed before sample removal for imaging. Fixed *Brucella* cells were vitrified on glow-discharged 200 mesh copper EM-grids with extra thick R2/2 holey carbon film (Quantifoil). Per grid, 3 μl of the sample was applied and automatically blotted and plunged into liquid ethane with the Leica EM GP plunge-freezer. Images were collected on a Talos L120C TEM (Thermo Fischer) using the Gatan cryo-TEM (626) holder. The images were acquired at a defocus between 8-10 μm, with a pixel size of 0.458 nm.

### Light microscopy images

Phase-contrast images of *B. abortus* and *B. ovis* cells from plates or liquid broth (plus or minus 1 mM IPTG) were collected using a Leica DM 5000B microscope with an HCX PL APO 63×/1.4 NA Ph3 objective. Images were acquired with a mounted Orca-ER digital camera (Hamamatsu) controlled by the Image-Pro software suite (Media Cybernetics). To prepare the different samples, cells were resuspended in PBS containing 4% formaldehyde.

### Agglutination assay

Agglutination assays were performed as previously described (8). The different *Brucella* strains (*B. ovis* and *B. abortus*) were harvested and resuspended in sterile PBS at OD600 ~ 0.5. One milliliter of each cell suspension was loaded in a spectrophotometer cuvette and mixed with 20 μl of wild-type *B. abortus*-infected mouse serum or with acriflavine (final concentration 5 mM) and OD was measured at 600 nm at time “0” and after 2 hours. As a control, 1 ml of each cell suspension was also kept in a spectrophotometer cuvette without serum or acriflavine.

### Alkaline phosphatase cell localization assay

To determine the cellular localization of EipB, we used a *B. ovis* strain transformed with the pSRK plasmid carrying *B. abortus eipB* C-terminally fused to *E. coli phoA*. Two versions of this plasmid were built: one carrying the full-length *eipB*, which expressed the protein with its signal peptide, and one carrying a short version of *eipB*, which expressed the protein lacking the signal peptide. Alkaline phosphatase assays were performed as previously described (8). Briefly, aliquots of overnight culture of *B. ovis* (grown in presence or absence of 1 mM IPTG) were mixed with 5-Bromo-4-chloro-3-indolyl phosphate (BCIP, final concentration 200 μM). After 2 hours of incubation, the color change was visually assessed and pictures were taken. The same experiment was performed with spent medium supernatants.

### Size exclusion chromatography

A DNA fragment corresponding to *B. abortus eipB* lacking the signal peptide (residues 31 - 280) was cloned into pET28a and transformed into the protein overexpression *E. coli* Rosetta (DE3) *pLysS* strain. Protein expression and purification was conducted using a Ni^2+^ affinity purification protocol as previously published (8). The purified protein was then dialyzed against a Tris-NaCl buffer (10 mM Tris (pH 7.4), 150 mM NaCl). EipB oligomeric state was analyzed by size exclusion chromatography as previously described (8). Briefly, after concentration, a protein sample (500 μl at 5 mg/ml) was injected onto a GE Healthcare Superdex 200 10/300 GL column (flow rate: 0.5 ml/min). Elution profile was measured at 280 nm and 500 μl fractions were collected during the run; the dialysis buffer described above was used for all runs. Protein standards (blue dextran / aldolase / conalbumin / ovalbumin) injected onto the column were used to construct a calibration curve to estimate the molecular weight of purified EipB.

### EipB expression, purification and crystallization

The DNA fragment corresponding to the *B. abortus* EipB protein (residues 31 - 280) was cloned into the pMCSG68 plasmid using a protocol previously published (8). For protein expression, an *E. coli* BL21-Gold(DE3) strain was used. Selenomethionine (Se-Met) protein expression and purification was performed as previously described (8). The purified protein was then dialyzed against 20 mM HEPES (pH 8), 250 mM NaCl, and 2 mM DTT buffer and its concentration was determined. The purified Se-Met EipB protein was concentrated to 160 mg/ml for crystallization. Initial crystallization screening was carried out using the sitting-drop, vapor-diffusion technique. After a week, EipB crystallized in the triclinic space group P1 from the condition #70 (F10) of the MCSG-2 crystallization kit, which contains 24% PEG1500 and 20% glycerol. Prior to flash freezing in liquid nitrogen, crystals were cryo-protected by briefly washing them in the crystallization solution containing 25% glycerol.

### Crystallographic data collection and data processing

Se-Met crystal diffraction was measured at a temperature of 100 K using a 2-second exposure/degree of rotation over 260°. Crystals diffracted to a resolution of 2.1 Å and the corresponding diffraction images were collected on the ADSC Q315r detector with an X-ray wavelength near the selenium edge of 12.66 keV (0.97929 Å) for SAD phasing at the 19-ID beamline (SBC-CAT, Advanced Photon Source, Argonne, Illinois). Diffraction data were processed using the HKL3000 suite (41). *B. abortus* EipB crystals were twinned and the data had to be reprocessed and scaled from the P21 space group to the lower symmetry space group P1 with the following cell dimensions: a= 47.36 Å, b= 69.24 Å, c= 83.24 Å, and **α**= 90.09°, **β**= 90.02°, **γ**= 78.66° (see Table S2). The structure was determined by SAD phasing using SHELX C/D/E, mlphare, and dm, and initial automatic protein model building with Buccaneer software, all implemented in the HKL3000 software package (41). The initial model was manually adjusted using COOT (42) and iteratively refined using COOT, PHENIX (43), and REFMAC (44); 5% of the total reflections was kept out of the refinement in both REFMAC and PHENIX throughout the refinement. The final structure converged to an R_work_ of 19.5% and R_free_ of 24.5% and includes four protein chains (A: 30-270, B: 31-271, C: 30-271, and D: 30-270), 9 ethylene glycol molecules, two glycerol molecules, and 129 ordered water molecules. The EipB protein contained three N-terminal residues (Ser-Asn-Ala) that remain from the cleaved tag. The stereochemistry of the structure was checked using PROCHECK (45), and the Ramachandran plot and was validated using the PDB validation server. Coordinates of EipB have been deposited in the PDB (PDB ID: 6NTR). Crystallographic data and refined model statistics are presented in Table S2. Diffraction images have been uploaded to the SBGrid diffraction data server (Data DOI: 10.15785/SBGRID/445).

### Disulfide bond reduction assays

DNA fragments corresponding to *B. abortus eipB* cysteine mutants (C69S, C278S, and C69S+C278S) and lacking the signal peptide (residues M1-A30) were cloned into pET28a and transformed into the protein overexpression *E. coli* Rosetta (DE3) *pLysS* strain. Protein expression and Ni^2^+ affinity purification were conducted using protocols previously published (8). Briefly, for each protein, a pellet corresponding to a 250 ml culture was resuspended in 1.5 ml of BugBuster Master Mix (MD Millipore) supplemented with 50 μl of DNAse I (5mg/ml). After 20 min on ice, cell debris was pelleted and the supernatant was mixed with 200 μl of Ni-NTA Superflow resin (Qiagen). Beads were washed with 8 ml of a 10 mM imidazole Tris-NaCl buffer (10 mM Tris (pH 7.4), 150 mM NaCl) and 5 ml of a 75 mM imidazole Tris-NaCl buffer. Proteins were eluted with 200 μl of a 500 mM imidazole Tris-NaCl buffer. 50 μl of each purified protein (at 0.5 mg/ml) were then mixed with 12.5 μl of a 4x protein loading dye containing or not 1 mM of DTT. Samples were boiled for 5 min and 10 μl were loaded on a 12% SDS-PAGE.

### Thermal shift protein stability assay

A thermal shift assay to assess protein stability was performed on 20 μl samples containing 25 μM of purified *B. abortus* EipB^WT^ or EipB^C69S+C278S^, 50x Sypro Orange (Invitrogen) and 2 mM DTT when needed. Each protein sample and solution was prepared with the same dialysis buffer (10 mM Tris (pH 7.4), 150 mM NaCl, 1 mM EDTA). Ninety-six-well plates (MicroAmp EnduratePlate Optical 96-well fast clear reaction plates; Applied Biosystems) were used and heated from 25 to 95°C with a ramp rate of 0.05°C/s and read by a thermocycler (QuantumStudio 5 real-time PCR system; Applied Biosystems - Thermo Fisher Scientific) using excitation and emission wavelengths of 470 ± 15 nm and 558 ± 11 nm, respectively. Protein Thermal Shift software v1.3 (Applied Biosystems - Thermo Fisher Scientific) was used for calculation of the first derivative of the curve to determine the melting temperature.

### Bacterial two-hybrid protein interaction assay

To assay EipB interaction with TtpA, we used a bacterial two-hybrid system (46). Briefly, a *B. abortus eipB* DNA fragment (lacking the signal peptide) was cloned into pKT25 vector and a *B. abortus ttpA* fragment (lacking the signal peptide) was cloned into pUT18 or pUT18C vectors. The different pUT18, pUT18C and pKT25 combinations were then co-transformed into a chemically competent *E. coli* reporter strain BTH101 and spotted on LB agar plates (ampicillin 100 μg/ml + kanamycin 50 μg/ml) supplemented with X-Gal (40 μg/ml).

### Pull-down assay between EipB and TtpA

To evaluate the interaction between *B. abortus* wild-type and cysteine mutant EipB and TtpA, the different genes were cloned into pET28a and pMAL-c2G expression plasmids and transformed in *E. coli* Rosetta (DE3) *pLysS* expression strain. The corresponding proteins (His6-EipB^WT^ or His6-EipB cysteine mutants, and MBP-TtpA) were overexpressed and purified using nickel affinity and amylose affinity gravity columns, respectively. Two milliliters of amylose resin were saturated with 10 ml of a clarified cell lysate corresponding to a 500 ml culture pellet of IPTG induced *Rosetta* pMAL-c2G-ttpA. Beads were thoroughly washed with 50 ml of a Tris-NaCl buffer (10 mM Tris (pH 7.4), 150 mM NaCl) and 200 μl of these beads were mixed with 500 μl of nickel purified EipB at ~0.5 mg/ml (see reference (8) for a detailed nickel-affinity purification protocol). After 30 min incubation in presence or absence of 1 mM DTT, the flow-through was saved and the beads were thoroughly washed with a Tris-NaCl buffer supplemented or not with 1 mM DTT. The protein was eluted with 200 μl of the same buffer containing 20 mM of maltose. The different protein samples (elutions and flow-throughs) were run on a 12% SDS-PAGE and Coomassie stained.

### Bioinformatics

Figures of the structures, structural alignments, electrostatic potential representations and root mean square deviation (rmsd) calculations were performed using PyMOL (PyMOL Molecular Graphics System, version 1.7.4; Schrödinger, LLC). Surface hydrophobicity was evaluated using the YRB python script (47). The XtalPred server (48) and Dali server (49) were used to identify proteins with the highest structural and sequence relatedness. The BLAST server (https://blast.ncbi.nlm.nih.gov/Blast.cgi) was used to identify homologs of *B. abortus* EipB in different taxa within the *Alphaproteobacteria*. The EipB weblogo was generated by aligning 447 DUF1849 protein sequences of *Alphaproteobacteria* retrieved from the EMBL-EBI website (https://www.ebi.ac.uk/interpro/entry/IPR015000/proteins-matched). Alignment was generated with Clustal Omega (https://www.ebi.ac.uk/Tools/msa/clustalo/). When necessary, the C-terminus of sequences were realigned by hand. The Clustal alignment file was converted to a fasta file using http://sequenceconversion.bugaco.com/converter/biology/sequences/clustal_to_fasta.php. This file was then submitted to skylign server (http://skylign.org/) to generate a weblogo. The alignment was processed with the following options: remove mostly-empty columns / alignment sequences are full length / score.

## Acknowledgments

We thank the members of the Crosson laboratory for helpful discussions. The authors wish to thank members of the SBC at Argonne National Laboratory for their help with data collection at the 19-ID beamline. This work was supported by National Institutes of Health Grants U19AI107792 and R01AI107159 to S.C.

## Author contributions

JH, JWW and SC contributed to the design and conceptualization of the study; JH, JWW, AF, DMC, JXC, EU, AB, LB, GB, YK and SC performed the experiments, acquired and analyzed the data; JH, JWW, AF and SC interpreted the data; JH and SC wrote the original draft of the manuscript.

