## Supplemental Figures and Tables for "*Brucella* periplasmic protein EipB is a molecular determinant of cell envelope integrity and virulence"

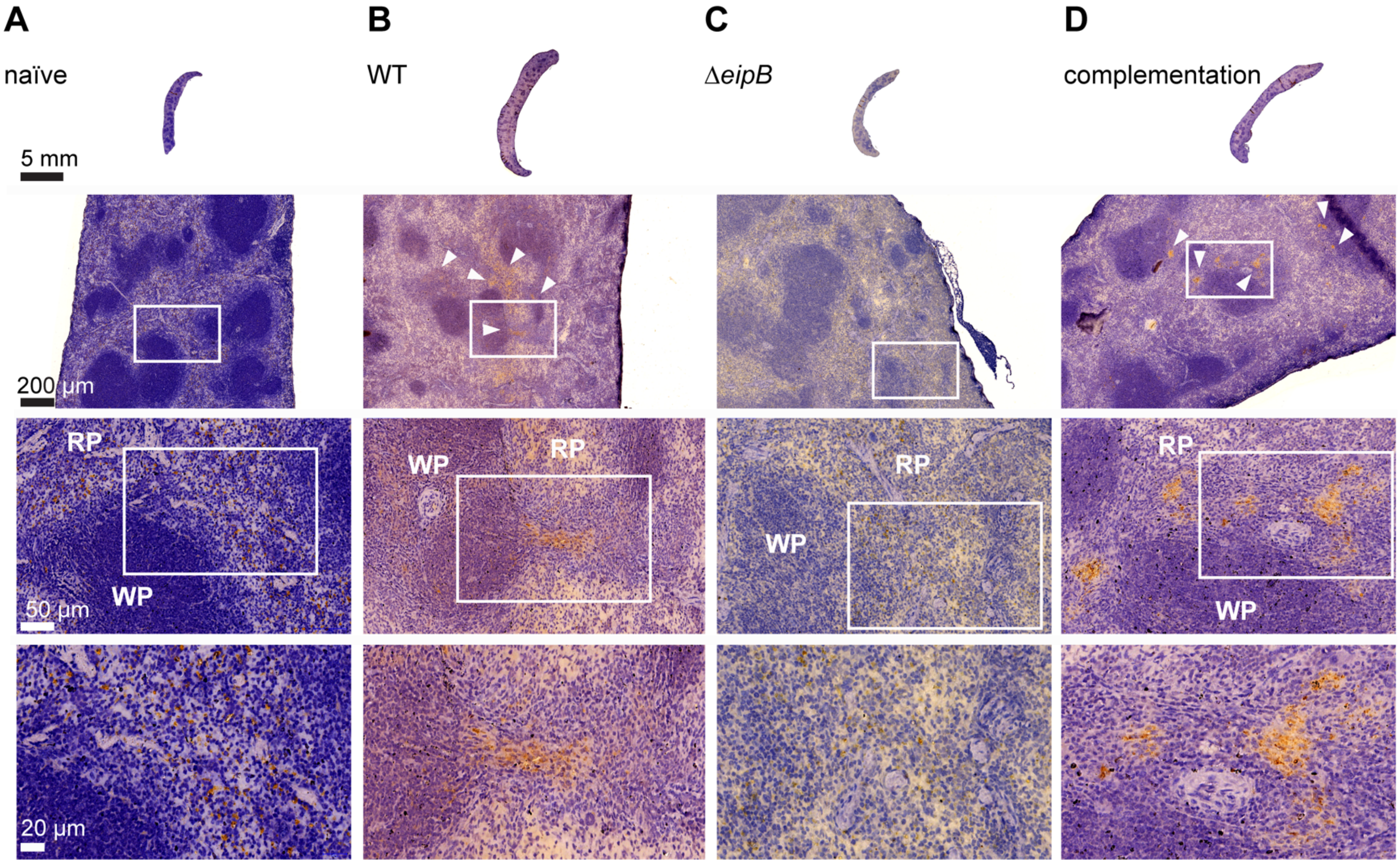


**Figure S1:** Histology of BALB/c mouse spleens at 8 weeks post-infection. Spleens (n= 1 per strain) were harvested, fixed, and stained with hematoxylin and eosin. Pictures of uninfected (A), wild-type (B), Δ*eipB* (C), or complementation (D) *B. abortus*-infected spleens were taken; white boxes represent specific regions enlarged in the images below. *Brucella* antigen was visualized by immunohistochemistry with an anti-*Brucella* antibody (brown regions, highlighted with white arrow heads). WP= white pulp, RP= red pulp.

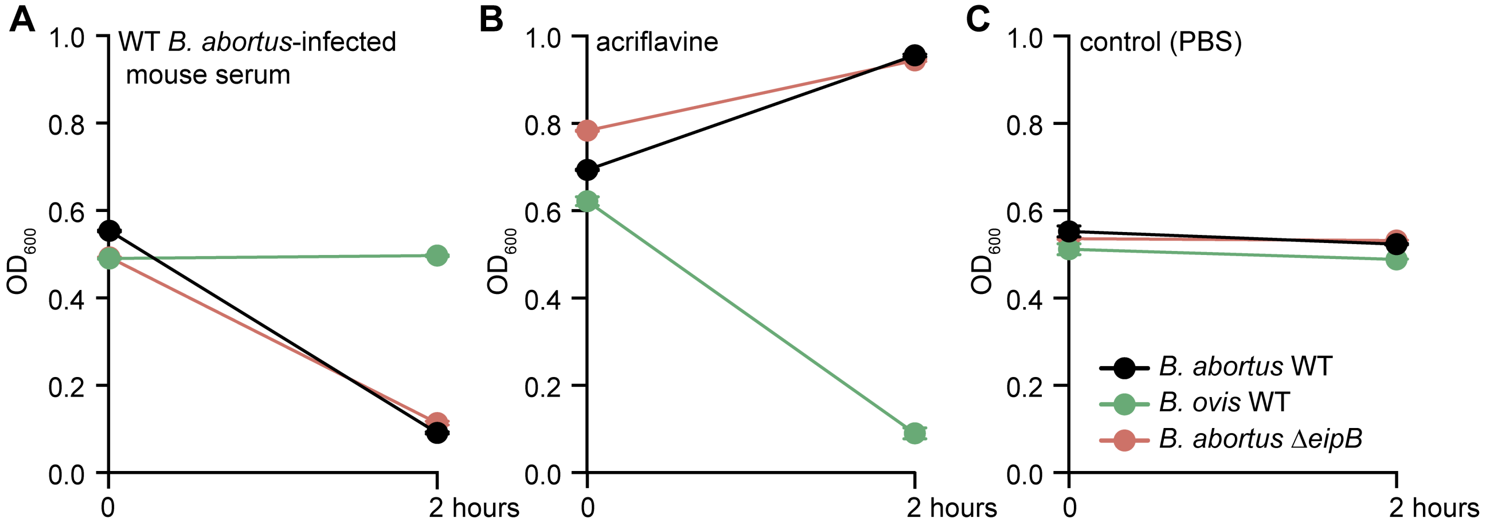


**Figure S2:** Deletion of *eipB* has no effect on the agglutination phenotype of *B. abortus*. The different *B. abortus* strains (wild-type in black, and Δ*eipB* in pink) and the wild-type *B. ovis* strain (in green) were incubated for 2 hours at room temperature in 1 ml of PBS supplemented with either (A) 20 µl of serum from a *B. abortus*-infected mouse, (B) 5 mM acriflavine (final concentration), or (C) no treatment. $OD_{600}$ was measured at the beginning and the end of the experiment. The starting OD used was ~0.5. This experiment was performed in triplicate; each data point is the mean ± the standard error of the mean.

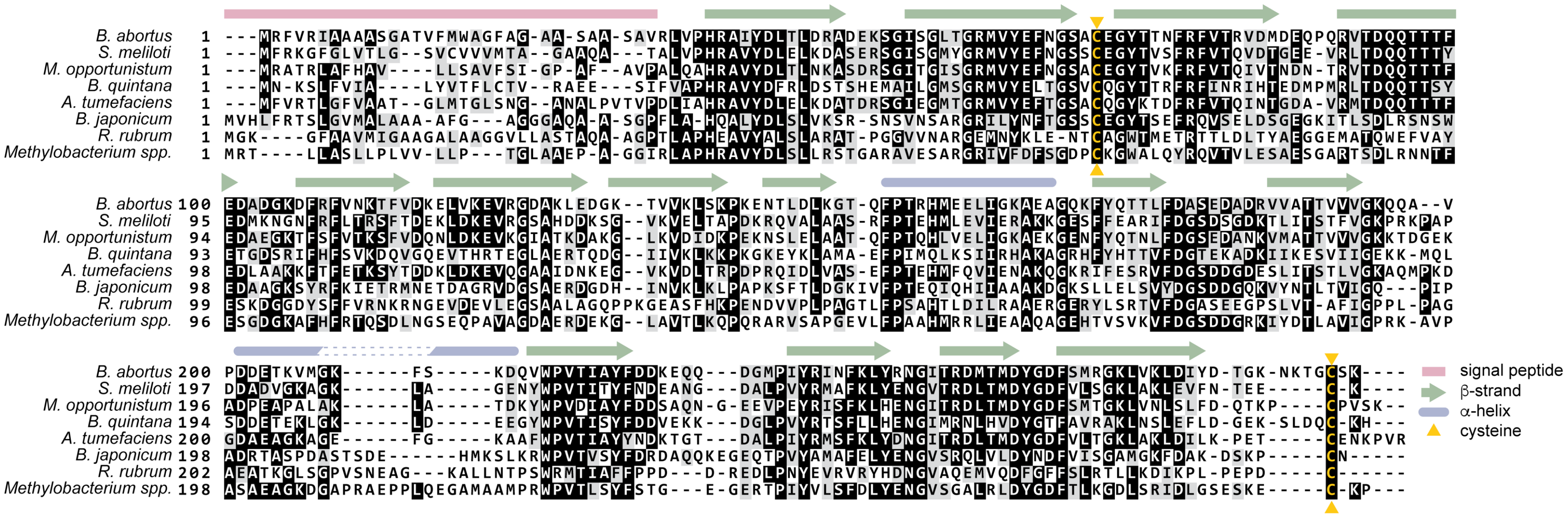

**Figure S3:** Amino acid sequence alignment of EipB (DUF1849) proteins of diverse *Alphaproteobacteria*: *B. abortus* (Bab1_1186), *Sinorhizobium meliloti* (SMc02102), *Mesorhizobium opportunistum* (Mesop_4249), *Bartonella quintana* (BQ07020), *Agrobacterium tumefaciens* (Ach5_12730), *Bradyrhizobium japonicum* (RN69_22205), *Rhodospirillum rubrum* (F11_10640), *Methylobacterium sp.* (M446_6683). Sequences corresponding to the peptide signal are delimited by a pink line. *B. abortus* EipB secondary structure is reported above the sequence alignment. β-strands are in green and α-helices are in violet. The conserved cysteines that form a disulfide bond are highlighted in yellow.

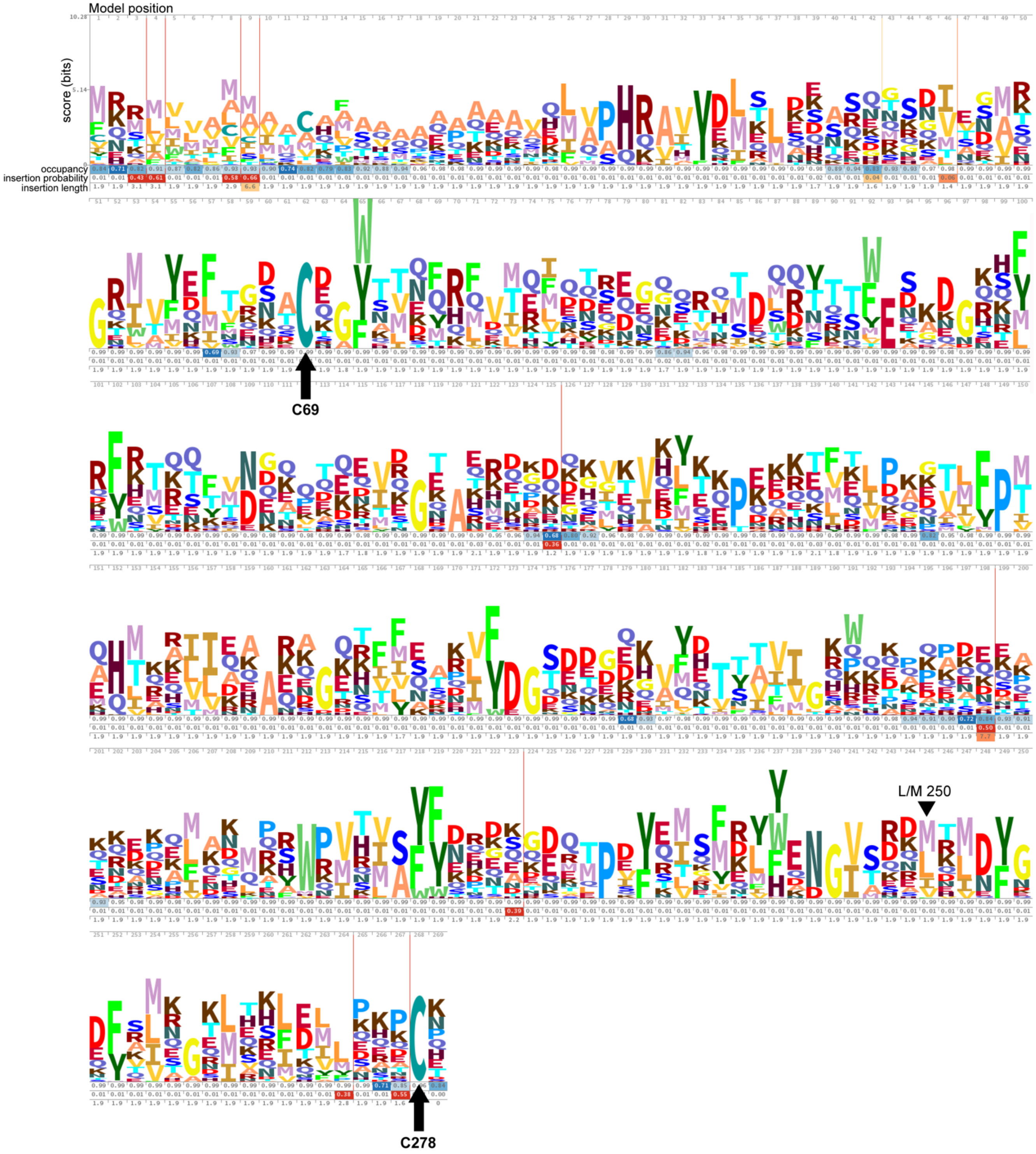


**Figure S4:** Weblogo of 447 DUF1849 protein sequences from diverse *Alphaproteobacteria*. Letter stacks and gap values depend on estimated distributions for each position. Those distributions were estimated by HMMER by applying sequence weights, absolute weights, and a Dirichlet mixture prior. In each position, letter height is the score of that letter at that position. Only positive-scoring letters are shown. Stack height has no inherent meaning. Positions of cysteines C69 and C278 in *B. abortus* EipB are reported on the model. These two cysteines are the most conserved residues in DUF1849 and form a disulfide bond. Position 250 is also highlighted with a black arrow.

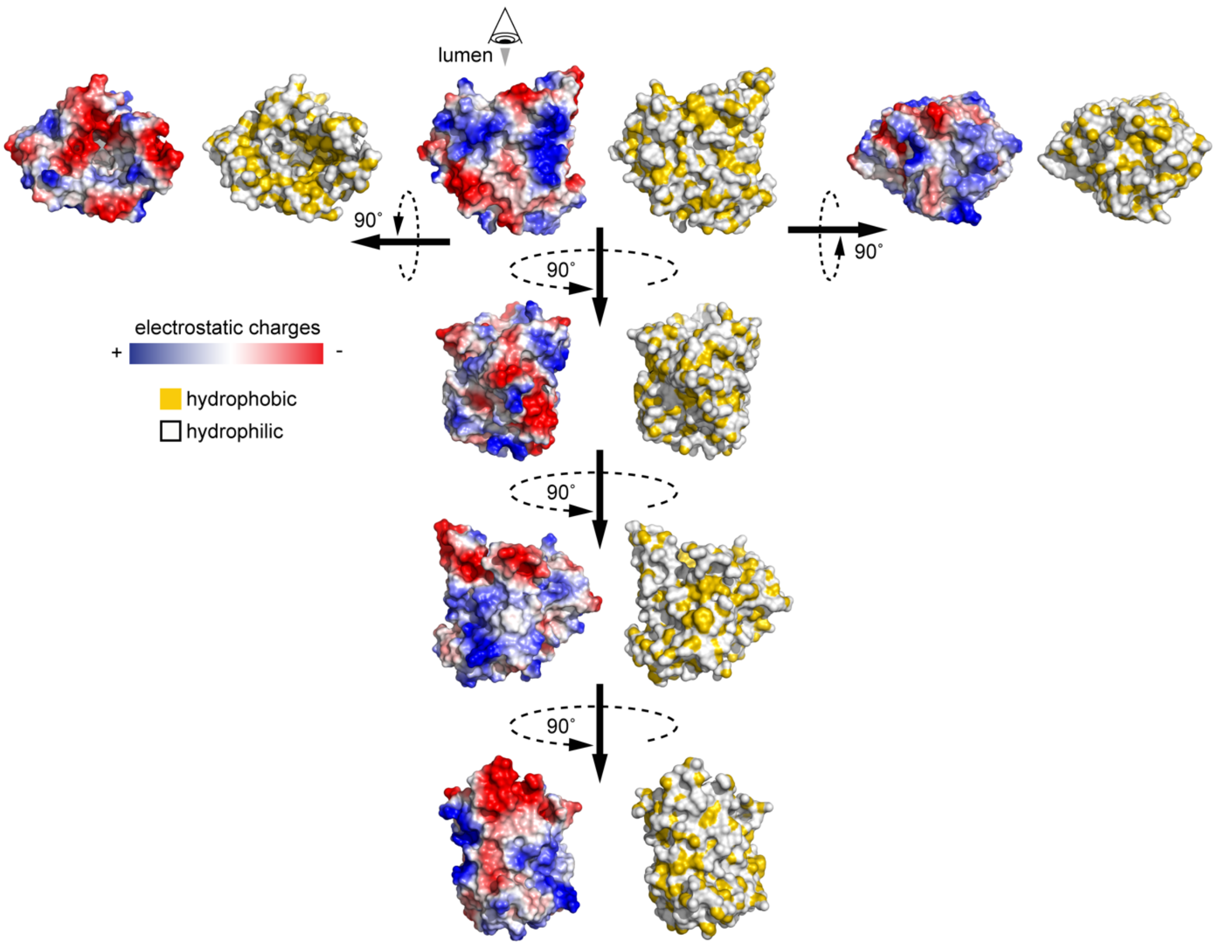


**Figure S5:** Surface representation of EipB in different orientations. Electrostatic potentials are mapped in blue for positive charges and in red for negative charges. Hydrophobicity is represented in yellow and hydrophilicity in white.

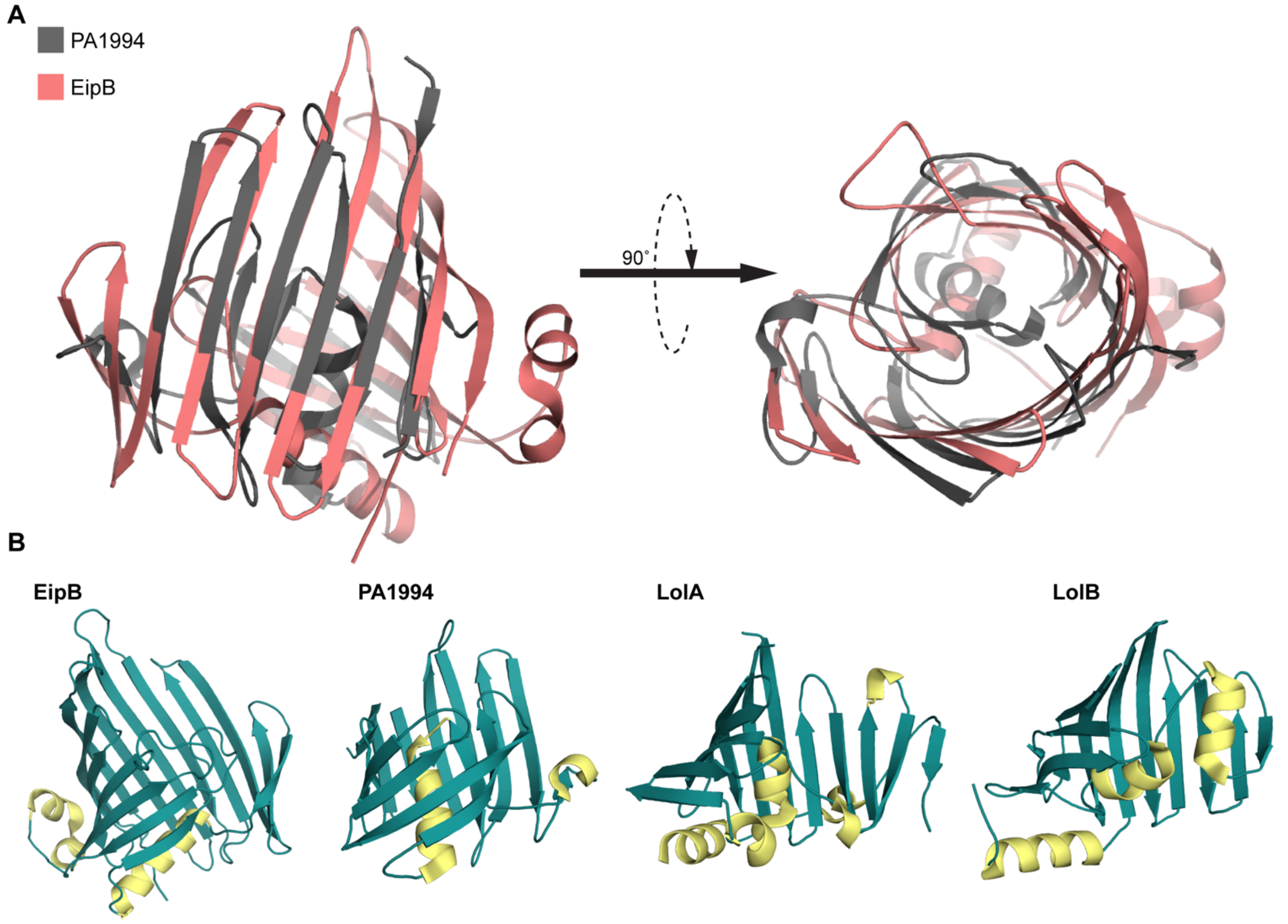


**Figure S6:** A) Structural alignment between EipB (PDB: 6NTR, in salmon), and *P. aeruginosa* P1994 (PDB: 2H1T, in grey). B) Structural feature comparison of EipB (PDB: 6NTR), *P. aeruginosa* PA1994 (PDB: 2H1T), *E. coli* LolA, (PDB: 1IWL), and *E. coli* LolB, (PDB: 1IWM). β-strands are in turquoise and α-helices are in yellow.

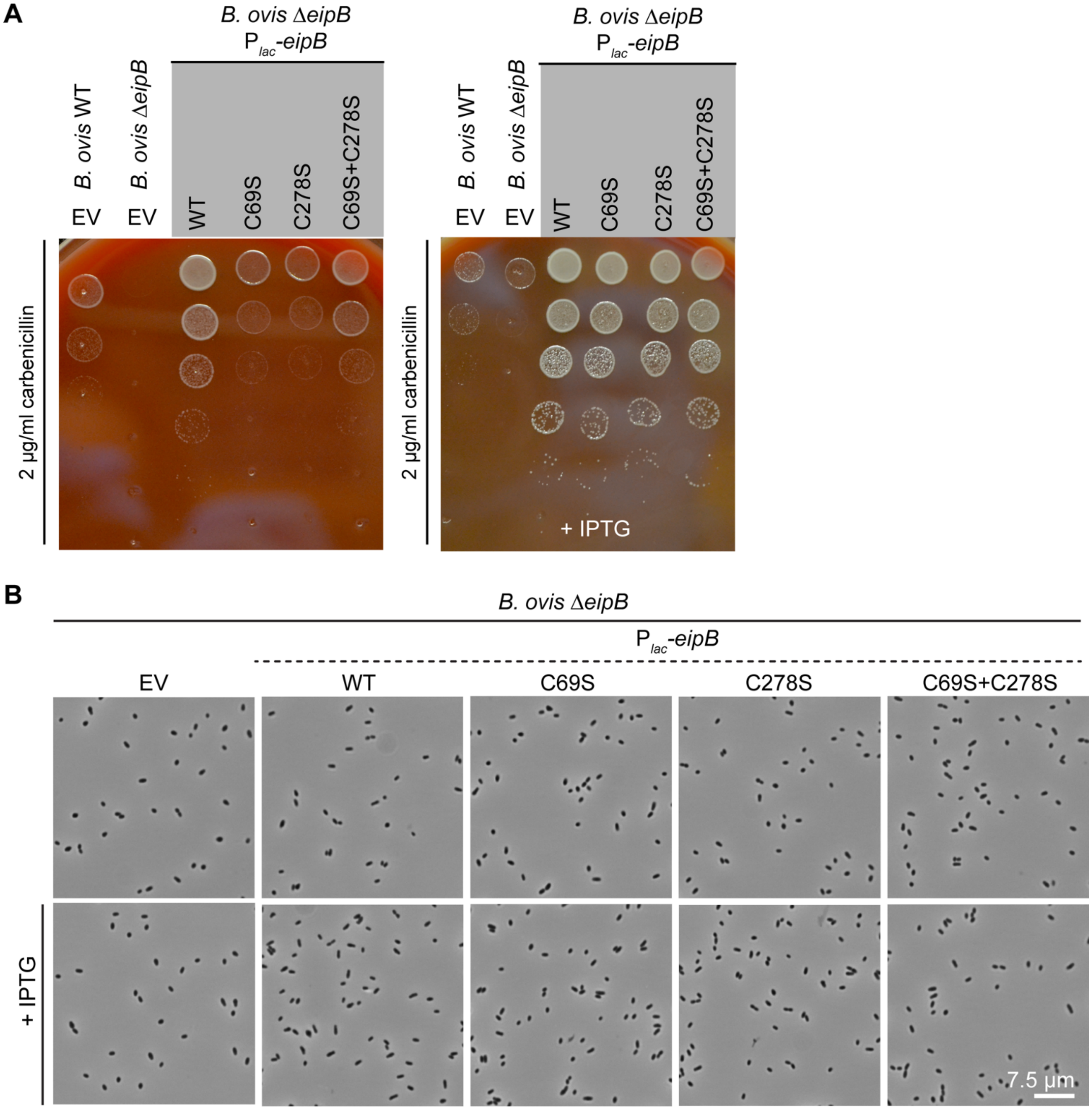


**Figure S7**: A) Stress assay performed on a SBA plate containing 2 µg/ml of carbenicillin. *B. ovis* Δ*eipB* expressing wild-type (WT) EipB systematically grew at 1 log higher dilution than the strains expressing EipB cysteine mutant; this difference is abolished in presence of 2 mM IPTG. Wild-type and Δ*eipB* *B. ovis* empty vector control strains are also shown. B) Phase contrast light micrographs of the *B. ovis* Δ*eipB* strains expressing wild-type (WT) EipB and the different EipB cysteine mutant versions from a *lac* promoter ($P_{lac}$). An empty vector (EV) strain was also used as control. Cells were harvested and resuspended in PBS + formaldehyde after 3 days of growth on SBA plates (+ 50 µg/ml kanamycin) plus or minus 2 mM IPTG.

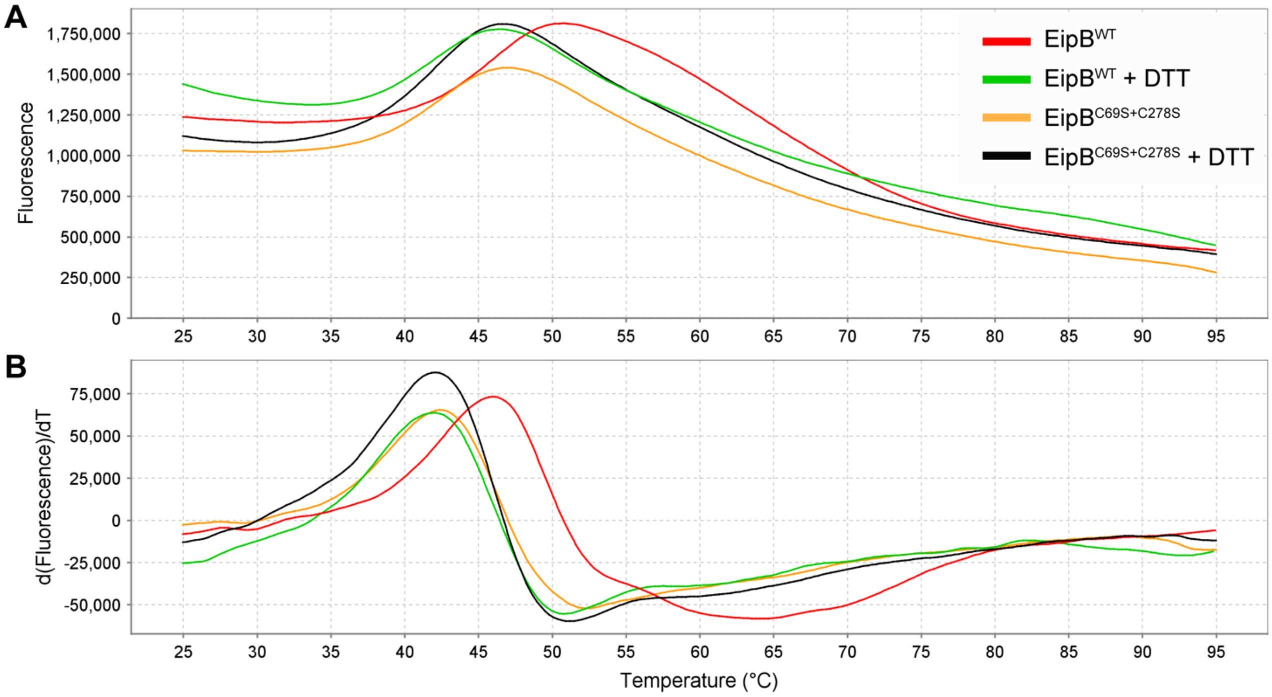


**Figure S8:** Thermal stability of wild-type EipB and EipB double cysteine mutant (that does not form a disulfide bond) in presence or absence of DTT. A) Representative melting profiles of purified EipB proteins (25 µM) in the presence or absence of 2 mM DTT. B) Derivative plots of the corresponding melt curve plots. Wild-type EipB has a temperature of melting of 45.89 ± 0.10°C in absence of DTT and 41.61 ± 0.12°C in presence of DTT. EipB$^{C69S+C278S}$ has a temperature of melting of 42.31 ± 0.13°C in absence of DTT and 42.35 ± 0.11°C in presence of DTT. For each protein, the melting temperature was independently measured twice in quadruplicate. The mean and the standard deviation were calculated.

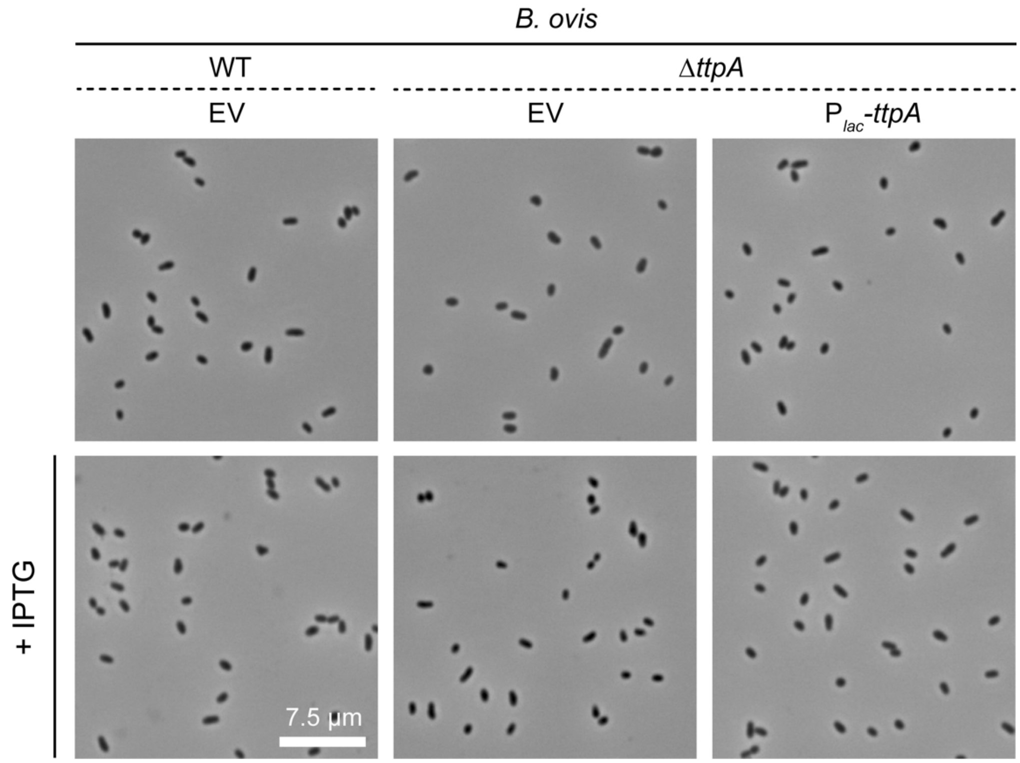


**Figure S9**: Phase contrast light micrographs of the *B. ovis* wild-type (WT) and Δ*ttpA* strains. The WT strain was transformed with the pSRK empty vector (EV) and the Δ*ttpA* strain was transformed with either the empty vector (EV) or with pSRK-*ttpA* ($P_{lac}$-*ttpA*). Cells were harvested and resuspended in PBS + formaldehyde after 3 days of growth on SBA plates (+ 50 µg/ml kanamycin) plus or minus 2 mM IPTG.

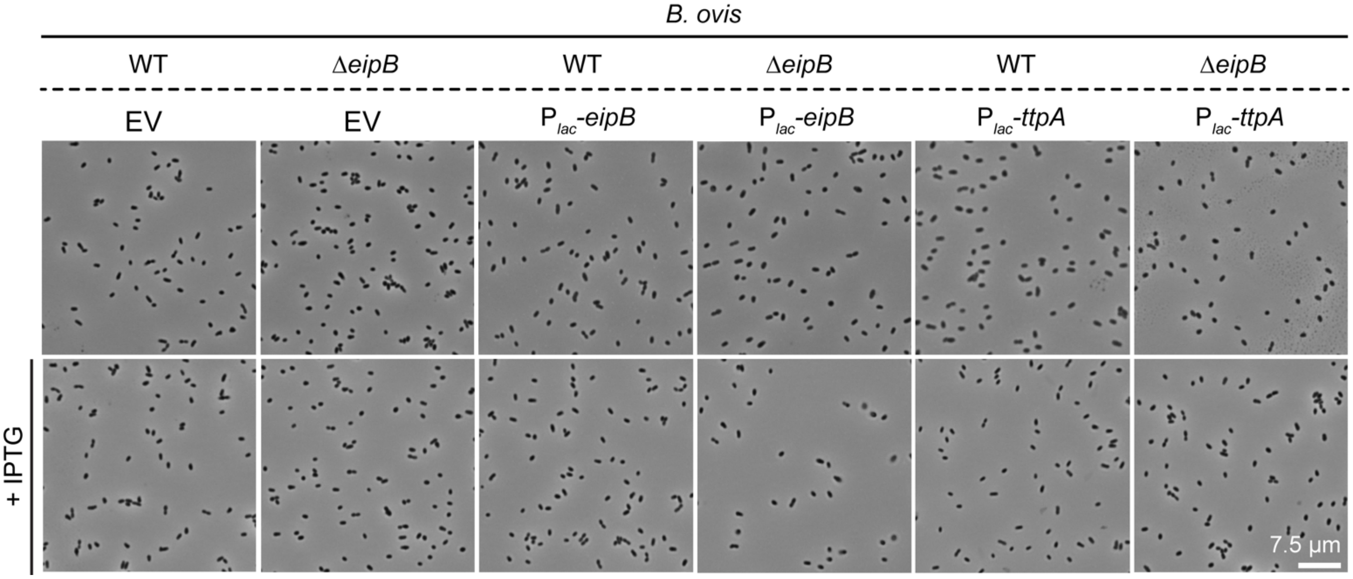


**Figure S10**: Phase contrast light micrographs of the *B. ovis* wild-type (WT) and Δ*eipB* strains expressing EipB or TtpA from a *lac* promoter ($P_{lac}$). Empty vector (EV) strains were also used as controls. Cells were harvested and resuspended in PBS + formaldehyde after 3 days of growth on SBA plates (+ 50 µg/ml kanamycin) plus or minus 2 mM IPTG.

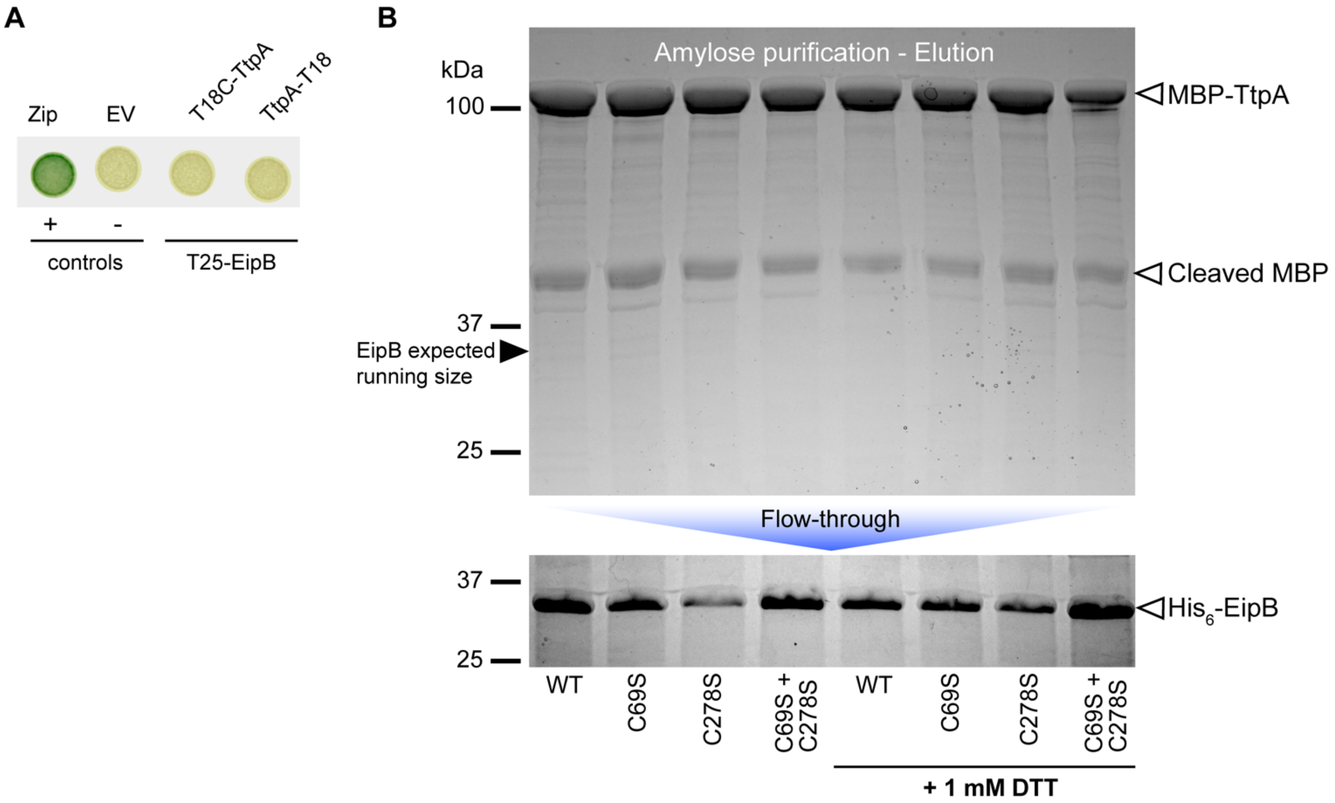


**Figure S11:** *Brucella* EipB-TtpA protein interaction assays. A) Bacterial two-hybrid (BTH) assay in which EipB is fused to the adenylate cyclase T25 fragment, and TtpA is N- or C-terminally fused to the T18 fragment. Leucine zipper (Zip) and empty vector (EV) are positive and negative controls, respectively. We found no evidence for interaction between EipB and TtpA by BTH. B) Affinity pull-down assay between His-tagged wild-type or cysteine (disulfide) mutant EipB (~31 kDa) and MBP-tagged TtpA (~109 kDa). Affinity purified $His_6$-EipB was loaded on a small column containing amylose resin that was saturated with MBP-TtpA. The flow-through was collected and the beads were thoroughly washed before elution. Flow-through and elution fractions were resolved by 12% SDS-PAGE. We found no evidence of EipB-TtpA interaction by affinity pull down.

**Table S1:** Summary of histopathology scoring of spleens from *B. abortus* WT- and Δ*eipB*-infected mice. Scores range from 0 to 3, and are based on masked evaluation of sectioned and stained spleen tissue. The presence of *Brucella* in tissue was confirmed by immunohistochemistry. 0= normal pathology (all naïve spleens scored 0 in all categories); 1= mild pathology, 2= moderate pathology, 3= severe pathology relative to uninfected (naïve) control.

| | **Naive** | **Wild-type** | **Δ*eipB*** | **Complementation** |
|---|---|---|---|---|
| **White pulp to red pulp ratio** | Normal (1:1) | Marked decrease, (2-3) | Minimal change, (0) | Mild decrease, (1) |
| **Average lymphoid follicles per field** | 12 | Decrease, (2) | Minimal change, (0-1) | Mild decrease, (1) |
| **Size of follicles** | Normal | Decrease, (2) | Minimal change, (0-1) | Mild decrease, (1) |
| **Marginal zone depletion** | Normal (intact) | Increase, (2) | Minimal to mild increase, (0-1) | Mild to moderate increase, (1-2) |
| **Extramedullary hematopoiesis** | Minimal | Moderate to marked increase, (2-3) | No change, (0) | Moderate increase, (2) |
| **Histiocytic proliferation** | Absent | Moderate to marked increase, (2-3) | Mild increase, (1) | Mild to moderate increase, (1-2) |
| **Granulomas** | Absent | Frequent, (3) | Rare, (1) | Occasional, (2) |
| **Brucella immunoreactivities** | Absent | Frequent, (2-3) | Rare, (1) | Occasional, (2) |

**Table S2:** Crystallographic data collection and refinement statistics. Statistics for the highest resolution shell are shown in parentheses.

| | **EipB** |
|---|---|
| **Wavelength (Å)** | 0.97929 |
| **Resolution range (Å)** | 35.23 – 2.10 (2.14-2.10) |
| **Space group** | P1 |
| **Unit cell** | a=47.36 Å, b=69.24 Å, c=83.24 Å, α=90.09°, β=90.02°, γ=78.66° |
| **# molecules in ASU** | 4 |
| **Unique reflections** | 57342 (2068) |
| **Multiplicity** | 2.8 (2.1) |
| **Completeness (%)** | 94.9 (68.1) |
| **Mean I/sigma(I)** | 20.0 (1.5) |
| **Wilson B-factor ($Å^2$)** | 53.6 |
| **R-merge** | 0.068 (0.390) |
| **cc1/2 (highest resolution shell)** | 0.876 |
| **Reflections used for R-free** | 2916 |
| **R-work** | 0.195 |
| **R-free** | 0.245 |
| **RMS(bonds)** | 0.012 |
| **RMS(angles)** | 1.474 |
| **Ramachandran favored (%)** | 96.5 |
| **Ramachandran outliers (%)** | 0.11 |
| **Clashscore** | 18.1 |
| **Average B-factor ($Å^2$)** | 70.1 |

**Table S3:** Transposon library statistics based on analysis of Tn-sequencing data using MapTnSeq.pl and DesignRandomPool.pl available at https://bitbucket.org/berkeleylab/feba. 150 bp single end read data that include the barcode and Tn-chromosome insertion junction are available through the NCBI sequence read archive (https://www.ncbi.nlm.nih.gov/sra) at accessions SRR7943723 (sequenced library constructed in wild type *B. abortus*), SRR7943724 (sequenced library constructed in wild type *B. ovis*), and SRR8322167 (sequenced library constructed in *B. abortus* Δ*eipB*).

| Library | Estimated number of Tn strains [a] | Unique Tn strain included [b] | Unique insertion sites[c] | Total TA sites[d] | Median Tn per gene | Median reads per gene |
|---|---|---|---|---|---|---|
| *B. abortus 2308* | $3.8 \times 10^6$ | 535,231 | 99,761 | 156,924 (78,462) | 86 | 1436 |
| *B. abortus 2308 ΔeipB* | $16.1 \times 10^6$ | 736,818 | 109,377 | 156,924 (78,462) | 122 | 6243 |
| *B. ovis* ATCC 25840 | $2.6 \times 10^6$ | 367,537 | 95,026 | 156,996 (78,498) | 61 | 1080 |

[a] Chao2 estimate of number of barcodes present in the library
[b] Number of unique barcodes passing mapping criteria (sequenced 5 or more times, mapped to unique site, perfect match)
[c] Counts hits on opposite strands as unique
[d] TA sites on forward and reverse complement (TA on forward only)

**Table S4:** (see Excel file Table S4) *B. abortus* WT, Δ*eipA*, and Δ*eipB* Tn-seq data.

**Table S5:** (see Excel file Table S5)**:** Primers and strains used in this study.
